# Extracellular release of damaged mitochondria induced by pre-hematopoietic stem cell transplant conditioning exacerbates graft-versus-host disease

**DOI:** 10.1101/2023.04.24.538141

**Authors:** Vijith Vijayan, Hao Yan, Juliane K. Lohmeyer, Kaylin A. Prentiss, Rachna V. Patil, Giulia Barbarito, Ivan Lopez, Aly Elezaby, Kolten Peterson, Jeanette Baker, Nicolai P. Ostberg, Alice Bertaina, Robert S. Negrin, Daria Mochly-Rosen, Kenneth Weinberg, Bereketeab Haileselassie

**Author notes:** Shared first authorship (Vijith Vijayan, Hao Yan). **Corresponding author:** Bereketeab Haileselassie MD MHS, Department of Pediatrics, Division of Critical Care Medicine, Stanford University School of Medicine, Stanford, CA.

## Abstract

Despite therapeutic advancements, GVHD is a major complication of HSCT. In current models of GVHD, tissue injury induced by cytotoxic conditioning regimens, along with translocation of microbes expressing Pathogen Associated Molecular Patterns (PAMPs), result in activation of host antigen-presenting cells (APC) to stimulate alloreactive donor T lymphocytes. Recent studies have demonstrated that in many pathologic states, tissue injury results in the release of mitochondria from the cytoplasm to the extracellular space. We hypothesized that extracellular mitochondria, which are related to archaebacteria, could also trigger GVHD by stimulation of host APC. We found that clinically relevant doses of radiation or busulfan induced extracellular release of mitochondria by various cell types, including cultured intestinal epithelial cells. Conditioning-mediated mitochondrial release was associated with mitochondrial damage and impaired quality control but did not affect the viability of the cells. Extracellular mitochondria directly stimulated host APCs to express higher levels of MHC-II, co-stimulatory CD86, and pro-inflammatory cytokines, resulting in increased donor T cell activation, and proliferation in mixed lymphocyte reactions. Analyses of plasma from both experimental mice and a cohort of children undergoing HSCT demonstrated that conditioning induced extracellular mitochondrial release *in vivo*. In mice undergoing MHC mismatched HSCT, administration of purified syngeneic extracellular mitochondria increased host APC activation and exacerbated GVHD. Our data suggests that pre-HSCT conditioning results in extracellular release of damaged mitochondria which increase alloreactivity and exacerbate GVHD. Therefore, decreasing the extracellular release of damaged mitochondria following conditioning could serve as a novel strategy for GVHD prevention.

## 1. Background

Allogeneic hematopoietic stem cell transplantation (HSCT) is a therapeutic strategy to treat various oncologic, hematologic, immune, and metabolic disorders by replacement of a patient’s blood-forming stem cells with those of a healthy donor ^(1)^. The broad application of allogeneic HSCT is limited by risks of several complications including graft-versus-host disease (GVHD), the leading cause of non-relapse mortality following HSCT^(2)^. GVHD occurs when alloreactive donor T lymphocytes recognize and direct an immune response against host cells ^(3,4)^. Current prevention and treatment strategies for GVHD rely on immunosuppressive medications that largely target the T cells which mediate the disease ^(5)^. However, therapies targeting the upstream mechanisms that initiate the GVHD immune response are limited.

Tissue damage from cytotoxic conditioning given before HSCT significantly contributes to the onset of GVHD. One explanation for the interrelationship between tissue damage and GVHD is the local or systemic release of microbe- and tissue-derived byproducts (pathogen associated molecular patterns (PAMPs) and damage associated molecular patterns (DAMPs) respectively), which prime host antigen presenting cells ^(6,7)^. Recently, extracellular mitochondria (exMito) have been identified as a link between tissue damage and inflammation in a variety of critical illnesses ^(8–12)^. Because of their archaebacterial ancestry, mitochondria retain molecular motifs which are^(8,11,12)^ recognized by pattern recognition receptors of the innate and adaptive immune system.

While extracellular release of mitochondrial content was previously thought to be a passive process secondary to cell death, recent discoveries suggest that it can be an alternative mechanism for mitochondrial quality control, especially in settings where the canonical mitochondrial quality control system is impaired ^(13,14)^. Our group, as well as others, have demonstrated that alterations in mitochondrial dynamics, in particular excessive mitochondrial fission, can mediate the extracellular release of damaged mitochondria following inflammatory insults ^(15,16)^. However, the impact of cytotoxic conditioning therapies on mitochondrial dynamics and the extracellular release of mitochondria have not been explored. In this article, we demonstrate that doses of irradiation or chemotherapy relevant to HSCT conditioning significantly increased mitochondrial fission and release of damaged mitochondria in various cell types, including intestinal epithelial cells (IEC). Furthermore, using an *in vivo* model of HSCT, we show that exMito increase the incidence and severity of GVHD. The pre-clinical findings are corroborated by an association between increased exMito content in circulation following conditioning and incidence of acute GVHD in a cohort of children undergoing HSCT.

## 2. Methods

### 2.1 Cell culture and murine strains

The human colon epithelial cell-line, Caco-2, human embryonic kidney (HEK293) and Human umbilical vein endothelial cells (HUVEC)(ATCC) were used to evaluate the impact of cytotoxic conditioning on mitochondrial morphology, function and exMito release. Caco-2 and HEK293 cells were grown in DMEM containing 4mM L-glutamine, 4.5g/L glucose and 1mM sodium pyruvate (Corning), 10% FBS (Genesee Scientific) and 1% penicillin and streptomycin. HUVECs were grown in Endothelial Cell Basal Medium containing endothelial Cell Growth supplement (Promocell). All cells were maintained at 37°C in an incubator with 5% CO2 and 100% humidity. All chemicals and reagents unless specified were purchased from Sigma Aldrich.

BALB/c and C57BL6/J Mice were acquired from Jackson laboratories. All animals had a at least 1-week acclimation period prior to initiation of experiments. All animal experiments were carried out under the protocol (#34455) approved by the Institutional Animal Care and Use Committee of Stanford University.

### 2.2 Cytotoxic pre-HSCT conditioning

Cell lines were treated with clinically relevant doses of pre-HSCT conditioning regimens (irradiation (5Gy and 10Gy) or busulfan (3μg/ml and 10 μg/ml)). Similarly, BALB/c mice were treated with varying doses of total body irradiation (TBI) (4.4 Gy and 8.8 Gy) or busulfan (20mg/kg/ day intraperitoneally for two doses) ^(17)^. C57BL6/J mice were treated with 10Gy of TBI. Following treatment, changes in mitochondrial morphology, function and extracellular release of mitochondria was evaluated as detailed in the supplemental methods.

### 2.3 Allogeneic HSCT model

BALB/c mice (8-12 weeks) were treated with fractionated total body irradiation [TBI] (2 doses of 4.4 Gy TBI on Day 0, eight hours apart) and were subsequently transplanted with 5×10^6^ T-cell depleted hematopoietic stem cells (HSC) from bone marrow alone or with 4×10^6^ splenocytes from C57BL6/J congenic mice expressing CD45.1+ allele to induce GVHD ^(18)^. Syngeneic damaged mitochondria along with bone marrow and splenocytes were administered to a subset of recipient BALB/c mice to determine the impact of circulating mitochondria on the incidence of GVHD. Syngeneic mitochondria were collected from heart, liver and kidney by differential centrifugation ^(15)^ and treated with 1mM H_2_0_2_ for 1 hour. Subsequently they were pelleted by centrifugation at 16000*g* for 20min, resuspended in MS buffer and the protein concentration was estimated by Bradford’s assay. Graded doses of damaged mitochondria (25 µg or 100 µg) were co-transplanted intravenously along with the bone marrow cells with or without donor splenocytes to determine the presence of a dose dependent response. Irradiated (8.8Gy) mice receiving either PBS, bone marrow, or bone marrow with mitochondria served as controls. Mice were monitored daily, and body weight and GVHD score were assessed twice weekly ^(19)^. A subset of animals was sacrificed at day 3 post transplantation and recipient (CD45.1 negative) cells from spleen and mesenteric lymph nodes were analyzed by flow cytometry for changes in MHC-II expression. The gating strategy used for flowcytometry analysis excluded doublets, Zombie Yellow^TM^ or Zombie Aqua^TM^(Biolegend) stained dead cells and CD45.1 stained donor cells.

### 2.4 Clinical cohort

Blood samples obtained from a prospective cohort of children undergoing HSCT at Lucile Packard Children’s Hospital (protocol ID: IRB 47220) were processed to deplete the platelets per previous protocols ^(20)^. 900 µL of MS buffer was added to 100 µL of plasma and centrifuged at 16000 g for 20 min. Supernatant was carefully removed and the mitochondrial pellet was resuspended in 100 µL of MS buffer. exMito abundance was quantified by flow cytometry using MTG staining (200 nM) as detailed in the supplemental methods. Abundance of exMito were evaluated pre and post conditioning and compared with incidence of acute GVHD.

### 2.5 Statistical analyses

Statistical analyses and graphing were performed using Prism 9.0 (Graphpad). Experimental results were evaluated based on the average of three or more independent experiments and calculated as mean values ± SD. Data distribution was evaluated with kernel density plots and the Shapiro-Wilk test for normality. For continuous variables, differences across groups were compared by one-way analysis of variance (ANOVA) or Kruskal-Wallis test with Bonferroni correction for multiple comparisons. Cox proportional-hazards model was used to assess survival differences across groups. The nominal p value of <0.05 was used as statistically significant threshold.

## 3. Results

### 3.1 Cytotoxic conditioning induces excessive mitochondrial fission and mitochondrial dysfunction in intestinal epithelial cells

To determine the impact of pre-HSCT irradiation conditioning on mitochondrial function, we treated intestinal epithelial cells (Caco-2) with escalating doses of irradiation (5Gy and10Gy) and characterized changes in bioenergetics, mROS and ATP production. Irradiated Caco-2 cells exhibited a significant decrease in mitochondrial oxygen consumption rate (Basal OCR: p=0.008; ATP-linked OCR: p=0.002; maximal OCR: p=0.03) (Fig. 1A) as well as basal glycolysis and glycolytic capacity (Supplemental Fig. S1). These impairments in cellular respiration were associated with a dose-dependent increase in mitochondrial oxidative stress (MitoSox MFI; 10Gy fold change relative to control = 1.41±0.04; p=0.0006) (Fig. 1B) and a decrease in complex I-dependent ATP production (10Gy fold change relative to control = 0.8 ± 0.09; p=0.005) (Fig. 1C).

**1).**
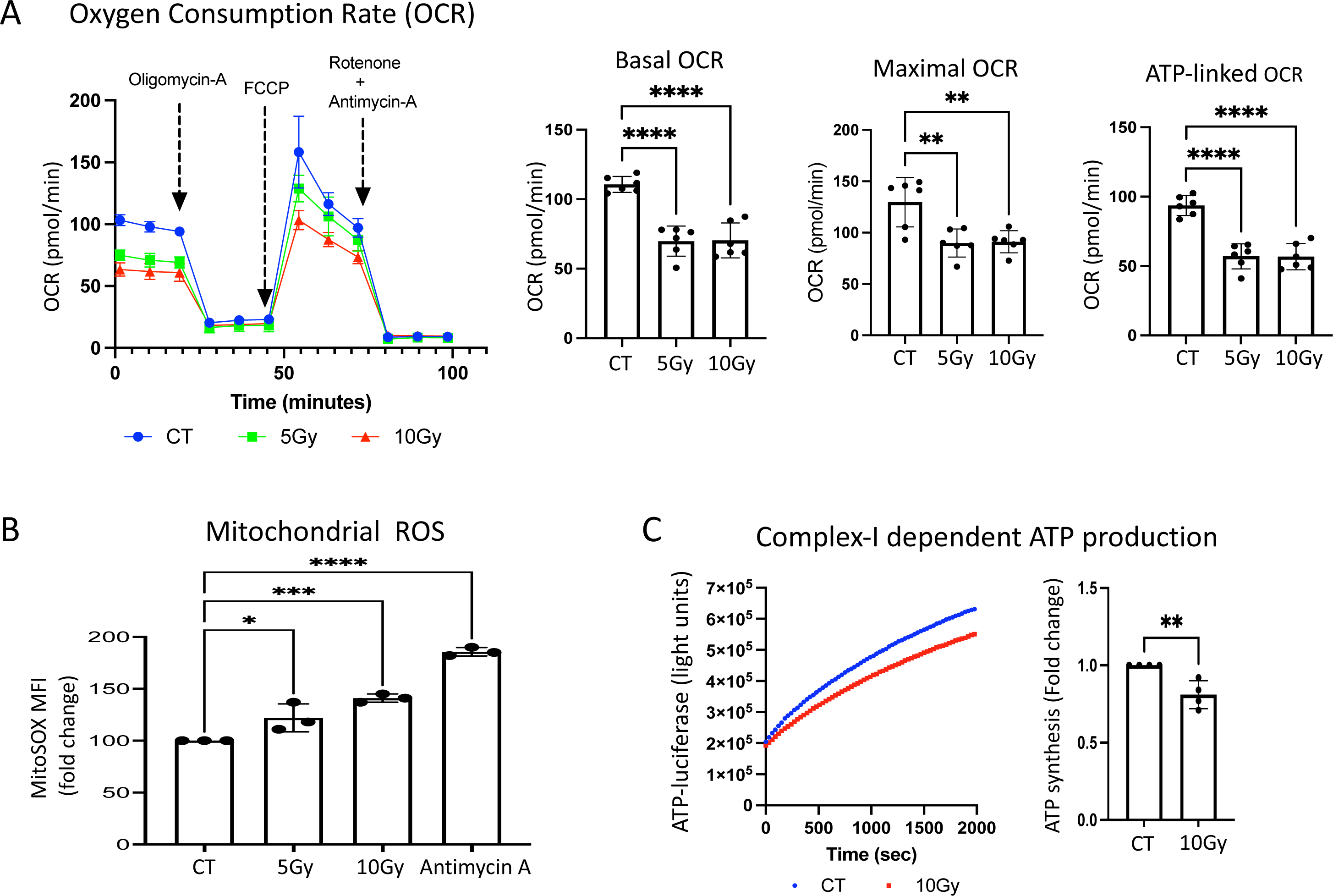
Irradiation leads to mitochondrial dysfunction in Caco-2 cells. Caco-2 cells were irradiated at the indicated doses, cultured for an additional 24 hours and subjected to various mitochondrial functional assays **(A)** A representative Seahorse plot of the Mito Stress assay and bar graphs of the respiratory parameters normalized within the experiment for difference in the total protein content between the groups (n=6). **(B)** The median fluorescence intensity of cells stained with MitoSOX^TM^ and analyzed by flow cytometry is shown as relative fold inductions in relation to control non-irradiated cells (CT) (n=3). Cells treated with 5 µM antimycin-A for 3 hours served as a positive control for the assay. **(C)** ATP production in 1µg of isolated mitochondria from non-irradiated (CT) and irradiated (10Gy) Caco-2 cells when provided with pyruvate and malate as substrates. A representative plot of luciferase activity recorded over time (left) and the ATP concentration calculated are shown as relative fold change in comparison to CT (n=4). (*p < 0.05, **p < 0.01, ***p < 0.001, ****p < 0.0001; n-number of independent experiments).

To validate our observations *in vivo*, we evaluated the effects of cytotoxic conditioning on mitochondrial dynamics and function by treating BALB/c mice with standard conditioning doses of irradiation (TBI 8.8Gy). Changes in mitochondrial architecture and function were then evaluated in intestinal cells as a target organ for GVHD. Electron microscopy of intestinal epithelial cells revealed mitochondrial damage with increased fission at 72 hours, followed by a loss of cristae density as well as outer mitochondrial membrane (OMM) integrity at 120 hours (Fig. 2A). Changes in mitochondrial morphology were quantified by a decrease in mitochondrial aspect-ratio (Control = 1.93±0.71 vs 8.8Gy = 0.8±0.6; p=0.001) and form factor (Control = 1.54±0.4 vs 8.8Gy = 0.51±0.32; p<0.001) (Supplemental Fig. S2A, S2B). Western blot analyses of isolated mitochondrial fractions from bowel showed an 8-fold increase in mitochondrial translocation of dynamin related protein-1 (Drp1; p=0.02), a key protein responsible for mitochondrial fission, in animals that received irradiation (Fig. 2B). Additionally, protein carbonylation assay of the mitochondrial fraction from small bowel showed a 1.8-fold increase in oxidative post-translational modifications (protein carbonylation; Control = 0.95±0.09 vs 8.8Gy = 1.8±0.47; p=0.01) following irradiation, suggesting an increase in oxidative stress (Fig. 2C).

**2).**
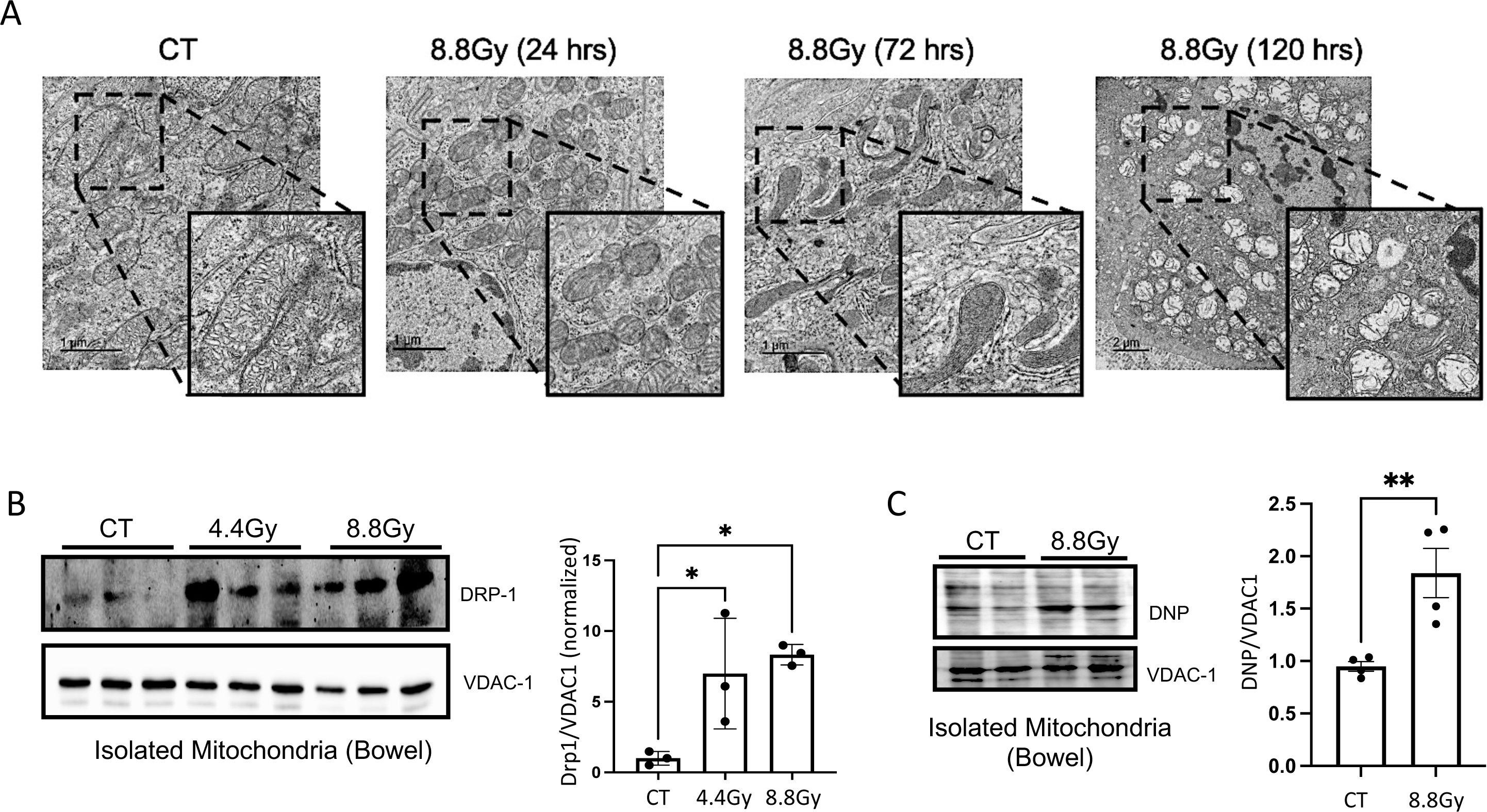
TBI induces mitochondrial fission and oxidative stress in the intestinal epithelium. Small bowel of BALB/c mice treated with TBI was resected at the indicated time points and evaluated for changes in mitochondrial architecture. **(A)** Representative electron micrograph for each time point is shown. **(B-C)** Enriched mitochondrial fractions isolated 24 h post TBI from small bowel of mice. **(B)** A representative Western blot probed with antibodies against DRP1 for mitochondrial fission, and VDAC-1 for total mitochondrial content (left) and densitometric protein quantification (n=3) (right) is shown. **(C)** A representative Western blot derivatized with dinitrophenylhydrazine (DNP) and probed with antibodies against DNP for protein carbonylation and VDAC-1 for total mitochondrial content (left) and densitometric protein quantification (n =3) (right) is shown. CT: control (non-irradiated) mice. (*p < 0.05, **p < 0.01; n – number of mice).

Under physiologic conditions, mitochondrial damage leads to the induction of mitophagy, the cellular quality control response for selective removal of damaged mitochondria. Analyses of intracellular mitochondrial content in total tissue lysates of small bowel showed no significant differences in the abundance of mitochondrial outer, inner membrane and matrix proteins between control and irradiated mice (Supplemental Fig. S3A, S3B,) indicating that dysfunctional mitochondria did not accumulate within the cells. However, western blot analyses of bowel tissue revealed that LC3-II levels (the lipidated form of LC3) in total tissue lysates (Supplemental Fig. S3A, S3B,) and the polyubiquitination status of mitochondrial proteins (Supplemental Fig. S3C, S3D) were not changed following irradiation, indicating no significant induction of either autophagy or mitophagy following irradiation. Absence of Pink-1 stabilization and Parkin translocation in the mitochondrial fractions further corroborated the lack of mitophagy induction by pre-HSCT radiation (Supplemental Fig. S3E, S3F). Altogether, these results suggest an alternate mechanism for clearance of dysfunctional mitochondria.

### 3.2 Cytotoxic conditioning induces extracellular release of damaged mitochondria

Besides mitophagy, extracellular release of mitochondria has been proposed to be an alternate mechanism of mitochondrial quality control following cellular stress ^(13,14)^. Because irradiation did not induce mitophagy, we hypothesized that pre-HSCT conditioning led to the extracellular release of damaged mitochondria. To test this hypothesis, we examined both mitochondrial release into the supernatant from cultured IEC and into plasma from mice treated with pre-HSCT irradiation. For the *in vitro* analyses, equal volumes of the cell culture supernatants from control and irradiated cells were subjected to differential centrifugation and the pellet was analyzed for mitochondrial presence by flow cytometry. Multiple mitochondrial markers including the mitochondria-specific stain MTG, mitochondrial membrane potential dye TMRM, and mitochondrial outer membrane protein (TOM20) were used to confirm that the yield from the differential centrifugation was indeed mitochondria (Supplemental Fig. S4A). In addition, exMito fractions were devoid of other associated organelles, including lysosomes, endoplasmic reticulum (ER) and proteasome as evaluated by western blot using antibodies against the respective organelle-specific markers (Supplemental Fig S4B)

Comparison of supernatants from control and irradiated Caco-2 cells revealed that irradiation caused approximately 50% increase in abundance of exMito (fold change in MTG positive events; 10Gy = 1.45±0.22; p=0.0002) (Fig. 3A). Similarly, Western blot analyses using antibodies against the complexes of the electron transport chain (ETC) indicated increased levels of mitochondrial proteins in the extracellular milieu of irradiated cells (Supplemental Fig. S5A). The increased release of mitochondria was independent of cell death, as measured by lack of significant difference in LDH release (Fig. 3B), suggesting that the expulsion of damaged mitochondria following irradiation was mediated by viable cells.

**3).**
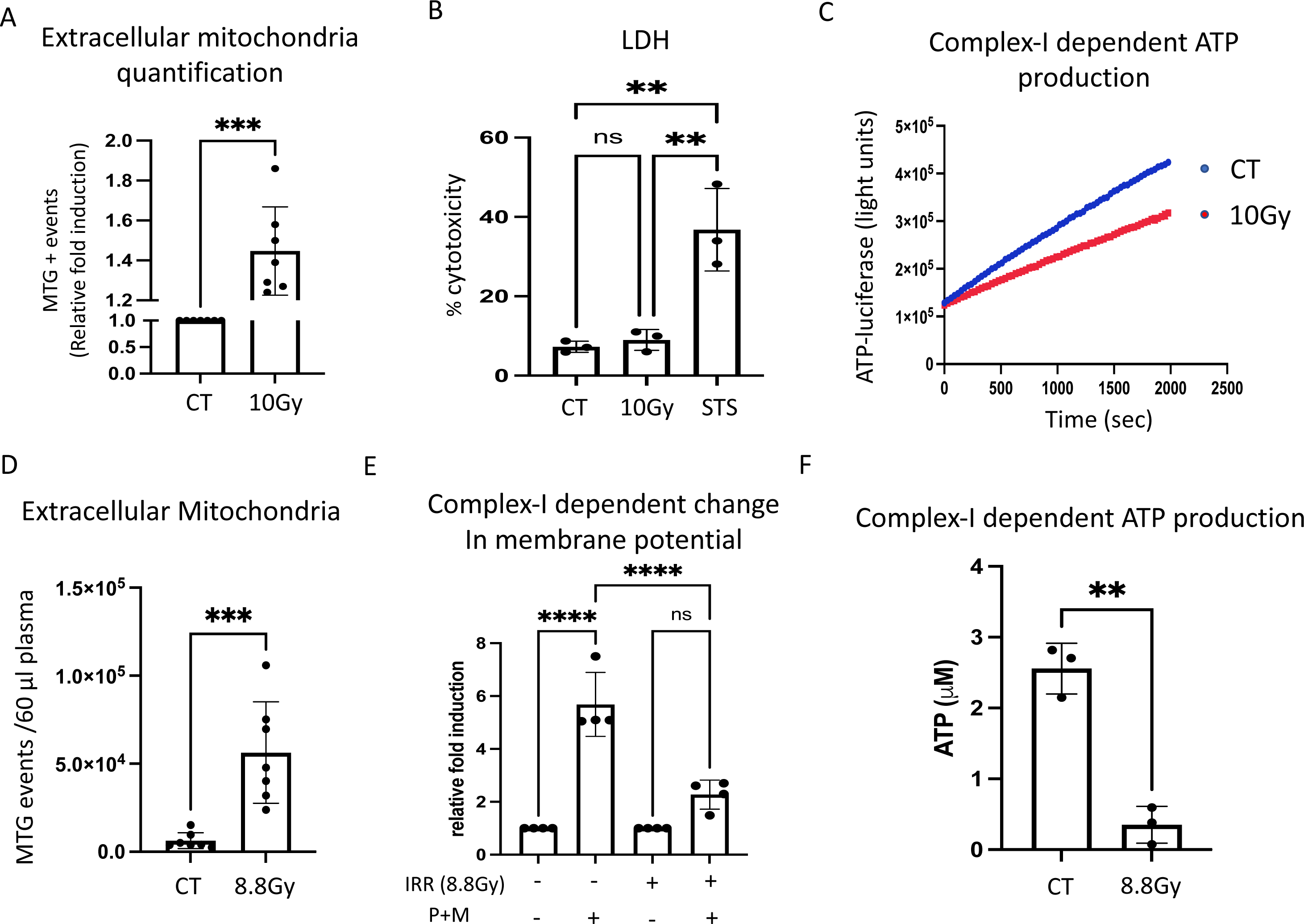
Irradiation leads to release of damaged exMito. exMito isolated from cell-culture supernatants of non-irradiated (CT) and irradiated (10Gy) Caco-2 cells was subjected to **(A)** MTG staining and analyzed by flow cytometry. The relative MTG positive events are shown as fold induction (n=7). **(B)** Cell culture supernatants from non-irradiated (CT) and irradiated (10Gy) Caco-2 cells cultured for 24 hours were subjected to LDH assay and the results are represented as % cytotoxicity. Staurosporine (STS) (1 µM) treatment for 12 hours was used as a positive control for the experiment (n=3). **(C)** complex-I dependent ATP-production in 1 µg of isolated mitochondria. A representative plot of luciferase activity recorded over time is shown. **(D)** exMito content in plasma of BALB/c mice 24 hours following TBI (8.8Gy) was stained with MTG was analyzed by flow cytometry. The number of MTG positive events detected in 60 µL of plasma is shown; CT-non-irradiated control mice (n=7). **(E)** exMito isolated from the plasma of BALB/C mice after 24 hours following TBI (8.8 Gy) was subjected to TMRM staining in the presence or absence of complex-I substrates (pyruvate and malate) and analyzed by flow cytometry. The median fluorescence intensity of TMRM calculated is represented as relative fold induction (n=4). **(F)** Mitochondria isolated from the plasma of BALB/C mice 24 hours following TBI (8.8Gy) or non-irradiated (CT) was subjected to complex-I-dependent ATP production (n=3). (*p < 0.05, **p < 0.01, ***p < 0.001, ****p < 0.0001; n-number of independent experiments).

When compared to isolated intracellular mitochondria (intMito), exMito were noted to be dysfunctional, represented by lower complex-I dependent ATP production (ATP (uM) per 1ug isolated mitochondria; intMito = 19.31± 3.74 uM vs exMito = 6.34 ± 3.5 uM, p=0.0006) (Supplemental Fig. S5B). Interestingly, the exMito released after irradiation demonstrated further impairment in ATP production (fold-change in ATP (uM) per 1ug isolated mitochondria relative to control; 10Gy = 0.65± 0.24, p=0.006) (Fig. 3C, Supplemental Fig. S5C) as well as oxidative stress (DNP/SHDB fold change; 10Gy = 2.1± 0.3 uM, p=0.0003) (Supplemental Fig. S5D).

To determine the impact of cytotoxic conditioning on exMito content *in vivo*, platelet-depleted plasma from control and irradiated BALB/c mice was collected. Flow cytometry of plasma with MTG stain revealed a significant increase in circulating cell-free mitochondria 24 hours following irradiation (MTG positive events/60 µL plasma; Control = 6,321±4,543 vs 8.8Gy = 56,349±28,824; p=0.0007), which persisted up to day 5 of analysis following conditioning (Fig. 3D, Supplemental Fig. S6A). The increase in cell-free mitochondria in plasma after irradiation was confirmed by western blot analysis of various mitochondrial proteins such as cytochrome c, and ATP5A1 (Supplemental Fig. S6B). Next, we evaluated the functional competency of exMito by testing their ability to hyperpolarize the mitochondrial membrane potential in response to complex-I substrates (pyruvate and malate (P+M)). Addition of complex-I substrates to exMito isolated from plasma of non-irradiated mice significantly increased the resting membrane potential, quantified by median fluorescence intensity of TMRM. However, exMito from plasma of irradiated mice exhibited diminished hyperpolarization (fold change; Control= 5.7±1.2 vs 8.8Gy = 2.3±0.55; p<0.0001) (Fig. 3E), indicating that exMito from irradiated mice have impairments in the production or maintenance of the proton motive force. This finding was further confirmed by reduced complex-I dependent ATP production from exMito of irradiated mice compared to nonirradiated controls (ATP (uM) per 1ug isolated mitochondria; Control = 2.6 ± 0.35 uM vs 8.8Gy = 0.35± 0.26 uM, p=0.001) (Fig. 3F and Supplemental Fig. S6C).

To confirm that this observation is not specific to BALB/c strain, we collected platelet depleted plasma from control and irradiated C57BL6/J mice and demonstrated significant increase in circulating cell-free mitochondria 24 hours following irradiation (MTG positive events/60 µL plasma; Control = 27,796 ±15,574 vs 10Gy = 51,638±14,632; p=0.008) (Supplemental Fig. S7A). Furthermore, to evaluate that these observations are also applicable to chemotherapy-based conditioning regimens, we treated Caco-2 cells with graded doses of busulfan and subsequently quantified exMito in the supernatant at 24 hours following treatment. Results demonstrated that busulfan treatment resulted in a 30% increase in exMito at 24 hours (p=0.001) (Supplemental Fig. S7B). Similarly, *in vivo*, mice treated with myeloablative doses of busulfan (20mg/kg IP x 2 doses)^(17)^ resulted in a significant increase in exMito 24 hours following treatment (MTG positive events/50 µL plasma; Control = 14,978±16,262 vs busulfan 20mg/kg IP = 30,639±11,541; p=0.03) (Supplemental Fig. S7C).

### 3.3 exMito increase activation of APC, induce T-cell proliferation, and increase the incidence and severity of GVHD in murine models

Activated host APCs play a crucial role in promoting the alloreactivity of donor T-cells after HSCT. To test the hypothesis that exMito stimulate host APCs to increase the allogenic response of donor T-cells, we used an *in vitro* mixed lymphocyte reaction (MLR) with responder T-cells from C57BL/6J mice and stimulator T-cell depleted splenocytes from BALB/c mice. To evaluate the impact of exMito, the stimulator T-cell depleted (BALB/c) splenocytes were incubated with and without exMito for 24 hours prior to MLR. Stimulation with exMito induced a dose dependent increase in the surface expression of the costimulatory molecule CD86 and MHC-II without change in MHC-I expression (Fig. 4A, 4B, Supplemental Fig. S8A), indicating an increased state of APC activation. Consistent with the altered expression of stimulatory molecules, the supernatants from the cultured splenocytes had significantly higher levels of cytokines and chemokines. Notably, cytokines relevant to GVHD that were induced by exMito included IFNγ, IL-10, IL-33, IL-6, MIP-2, IL-17, IL2Ra ^(21–28)^ (Fig. 4C, Table S1). Further evaluation of APC subpopulations demonstrated that B-cells and monocyte/macrophages have the most significant increase in the surface expression of the costimulatory molecule CD86 and MHC-II following exposure to exMito while dendritic cells have minimal response (Fig. 4D, 4E, Supplemental Fig. S8B). Together, the data demonstrate that exMito activate APC to express both surface and secreted proteins that are predicted to lead to greater T cell activation, differentiation, and expansion.

**4).**
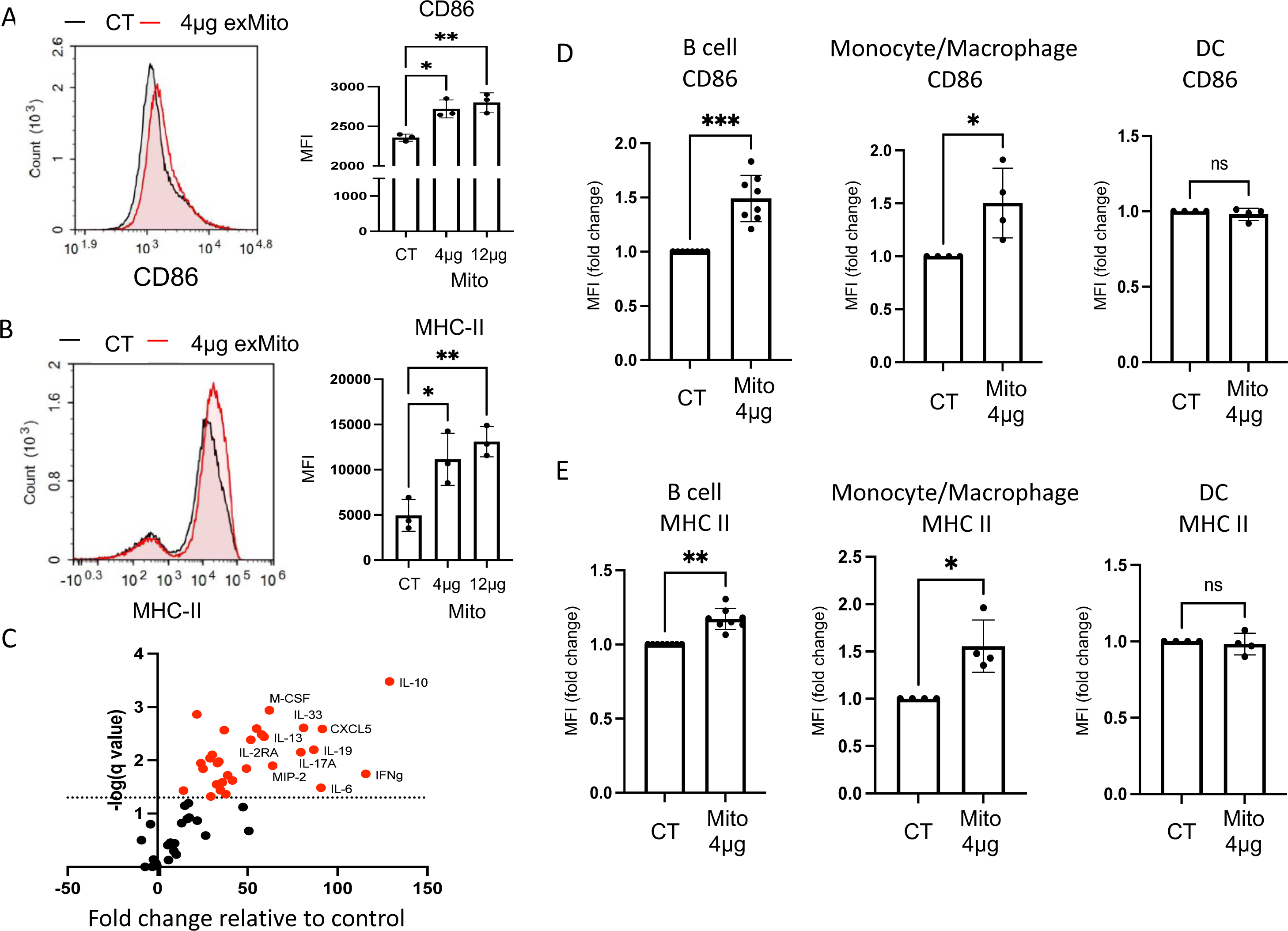
exMito activates antigen presenting cells *in vitro*. T-cell depleted BALB/c splenocytes were treated with 4 µg or 12 µg of mitochondria for 24hours. Cells were stained with **(A)** CD86 and **(B)** MHC-II antibodies respectively. A representative histogram and the calculated median fluorescence intensity (MFI) analyzed by flow cytometry is shown (n=3)**. (C)** Changes in cytokine levels in cell culture supernatants between control and 4 µg of mitochondria treated splenocytes is shown in volcano plot. Fold change relative to control demonstrated on x axis and inverse log p value demonstrated in y. Chemokines and cytokines highlighted in red were noted to be significantly different (n = 6). **(D-E)** The impact of exMito on different APC subpopulation including B cell, macrophages/monocytes and dendritic cells were further characterized (gating protocol described in supplemental Fig. S8B). Expression of **(D)** CD86 and **(E)** MHC-II quantified as fold change in median fluorescence intensity (MFI) is shown (n = 4-8). (*p < 0.05, **p < 0.01, ***p < 0.001; n-number of independent experiments).

To test the functional consequences of APC activation by exMito on interacting T lymphocytes, we evaluated the donor T cell activation and proliferation in the aforementioned MLR experiments. exMito treated BALB/c stimulators significantly increased the surface expression of the early activation marker CD69 on the allogeneic responder T-cells (Fig. 5A). Similarly, exMito-treated stimulator splenocytes increased proliferation of responder T-cells in comparison to untreated stimulators (Fig. 5B). Further characterization of the T-cell populations at 48 hours of MLR demonstrated that exMito treated splenocytes increased CD25 expression on both CD4^+^ and CD8^+^ T-cells (Fig. 5C). However, no significant differences were observed at 48 hours between the groups in the differentiation of CD4^+^ and CD8^+^ T-cells to central memory and effector cell populations as analyzed by the expression of CD44 and CD62L (Fig. 5D). Additionally, exMito-treated stimulators did not alter the abundance of regulatory T-cells in culture (Supplemental Fig. S9) indicating that the increased proliferation of conventional T-cells was not due to a reduction in regulatory T-cell abundance. Altogether, these data indicates that exMito stimulation of recipient APC leads to increased activation donor T-cells *in vitro*, an effect that was predicted to exacerbate GVHD *in vivo*.

**5).**
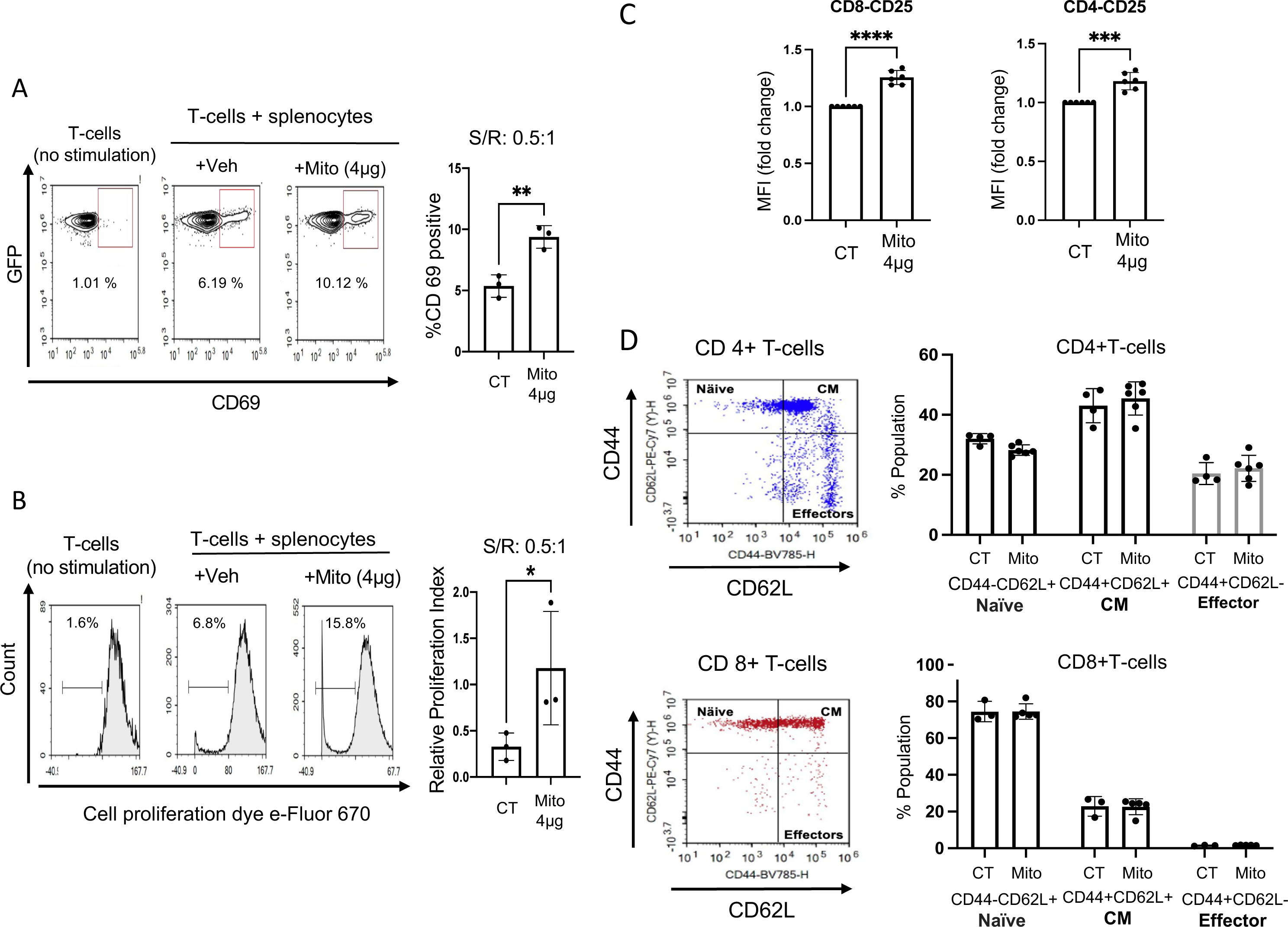
exMito treated APCs increase allogeneic T-cell activation and proliferation in mixed lymphocyte reaction. *In vitro* MLR between C57BL/6J responder T-cells and BALB/c T-cell depleted splenocytes was set up as described in the Methods section. **(A)** Cells were stained with a CD69 antibody and representative dot plots for the expression of CD69 on GFP-expressing responder T-cells after 48 hours of MLR (S/R: 0.5 to 1) (left) and the mean percentage CD69 positive T-cells (right) are shown. **(B)** Representative histograms after 120 hours of MLR with the indicated treatments showing the percentage of proliferate responder T-cells (left) and relative proliferation index (RPI) calculated using the equation: RPI = (stimulated T-cells with or without mitochondrial treatment - non stimulated T-cells)/ stimulated T-cells without mitochondrial treatment (right) is shown (n=3). **(C)** CD25 expression on CD4 and CD8 positive allogenic T-cells after 48 hours presented as MFI relative fold change (n=6). (D) percent of naïve (CD44-CD62L+), central memory (CM) (CD44+CD62L+) and effector (CD44+CD62L-) CD4+ and CD8+ T-cells after 48 hours in a MLR (n=4-6). (*p < 0.05, **p < 0.01, ***p < 0.001; n-number of independent experiments).

To validate our *in vitro* findings and determine the impact of exMito on incidence and severity of acute GVHD (aGVHD), we utilized a well-characterized aGVHD model ^(18)^ in which T-cell depleted BM cells (5×10^6^ cells/mouse) and splenocytes (4×10^6^ cells/mouse) from C57BL6/J donors were transplanted into irradiated BALB/c recipient mice, which received increasing doses of damaged mitochondria (vehicle control, 25, 100 µg) (Fig. 6A). Dosing with isolated exMito in addition to TBI significantly increased the abundance of circulating cell free mitochondria in comparison to mice that received TBI alone (Supplemental Fig. S10). Flow cytometric analysis at day 3 post-transplantation revealed an increased expression of MHC-II in recipient cells isolated from spleen and mesenteric lymph nodes (MLN) of mice which were treated with mitochondria, confirming the observed effect of exMito on isolated splenocytes *in vitro* (Fig 6B). BALB/c mice that were transplanted with HSCs and splenocytes had a median survival of 31 days, which was worsened when co-treated with exMito, (median survival; BM + splenocyte + mitochondria (25ug) = 21days, p=0.0042; BM + splenocyte + mitochondria (100 µg) = 14 days, p= 0.0014) (Figs. 6C, 6D). To confirm that the accelerated mortality following treatment with exMito is GVHD-specific, we transplanted irradiated BALB/c recipient mice with T cell-depleted BM cells (5×10^6^ cells/mouse) from C57BL6/J donors with and without damaged mitochondria (100 ug), but without donor splenocytes. In the absence of donor splenocytes, exMito did not significantly increase mortality (Fig. 6C). These findings along with the MLR results (Fig. 5A-C) suggest that the impact of damaged exMito is dependent on the presence of alloreactive T-cells which are necessary for GVHD.

**6).**
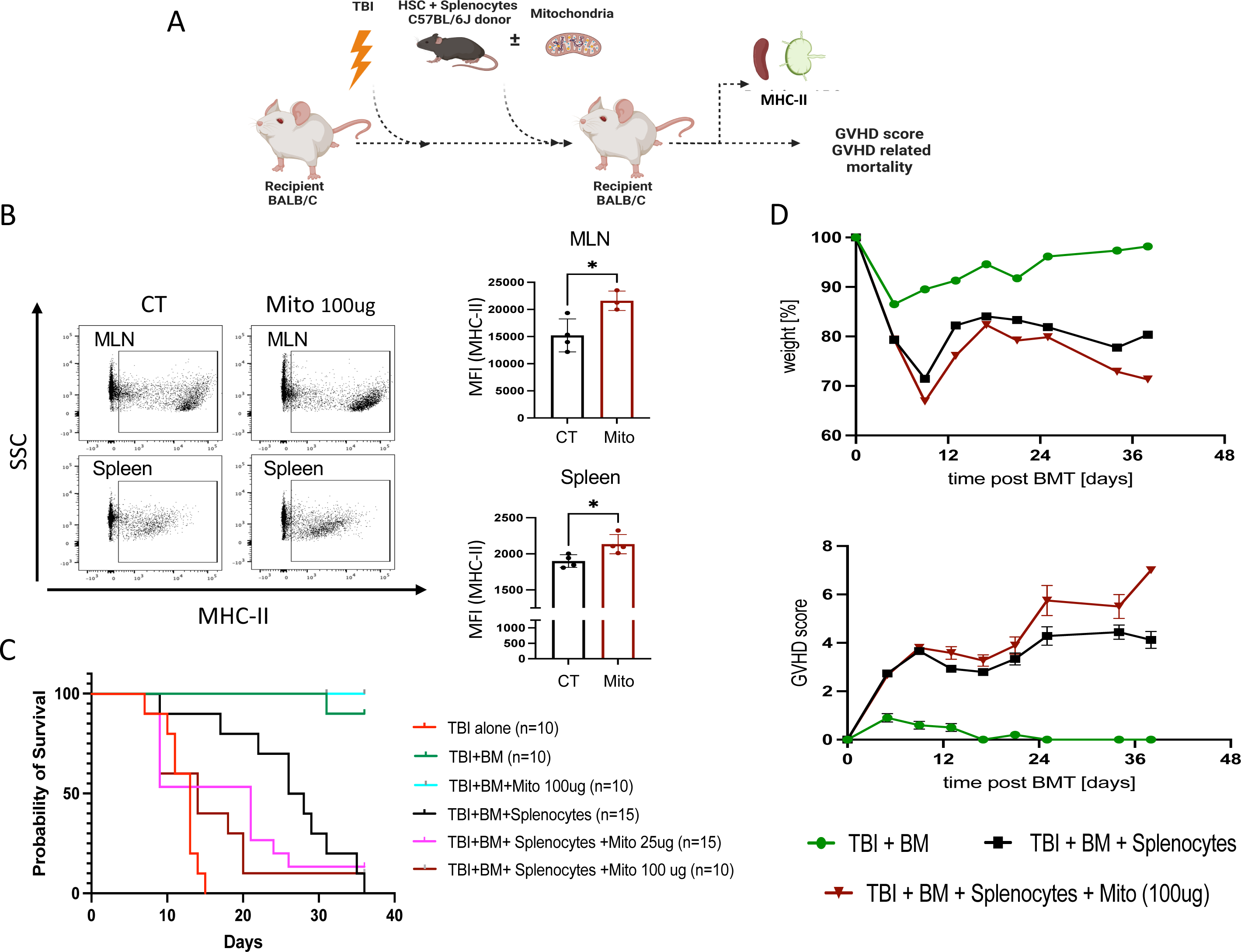
exMito aggravate GVHD-induced mortality. Hematopoietic stem cells (HSC) from bone marrow (5×10^6^ cells) and splenocyte (4×10^6^ cells) from C57BL6/J congenic mice expressing CD45.1+ allele were transplanted into BALB/c CD45.2+ mice following TBI conditioning to model acute GVHD. At the time of transplantation, the indicated doses of mitochondria were given along with the bone marrow. **(A)** Graphical representation of the experimental setup**. (B)** Isolated recipient CD45.2+ cells from spleen and mesenteric lymph nodes (MLN) were analyzed for the expression of MHC-II. Donor cells were excluded from analysis using an antibody against CD45.1 Representative dot plots (left) and MFI (right) shown as a bar graphs. (Mito: mitochondria-100 µg) **(C)** survival curve **(D)** illness severity including weight loss, and GVHD scores. (*p < 0.05, **p < 0.01, ***p < 0.001, ****p < 0.0001; n-number of animals).

### 3.4 Association of increased circulating mitochondria with conditioning and GVHD in children undergoing HSCT

To validate the clinical relevance of our *in vitro* and *in vivo* findings, we evaluated changes in exMito content from platelet-depleted plasma samples of pediatric patients undergoing HSCT. Transplant indications for the cohort consisted of hematologic cancers, nonhematologic cancers and bone marrow failure syndromes requiring diverse therapies prior to initiation of HSCT (Table 1). When stratified by incidence of GVHD, we did not observe any significant differences in patient demographics, indication for transplant, HLA molecular typing or GVHD prophylaxis (Table 1). We observed an overall increase in exMito abundance following conditioning, which was most pronounced in the subset of patients who received TBI as a component of their conditioning regimen, (MTG positive events/60ul plasma fold change from baseline; day 0 = 4.03 ± 1.5 p= 0.047, day 7= 2.43± 0.71 p= 0.043, day 14= 3.7± 1.3 p= 0.016), (Fig. 7A). Compared to those who did not develop GVHD, the patients who later developed GVHD had a 6-fold higher level of exMito content on Day 0 (day of transplantation) (MTG positive events/60ul plasma; no-GVHD = 6,023 ± 13,239 vs GVHD = 56,222 ± 61,131, p=0.04) (Fig. 7B).

**Table 1:**
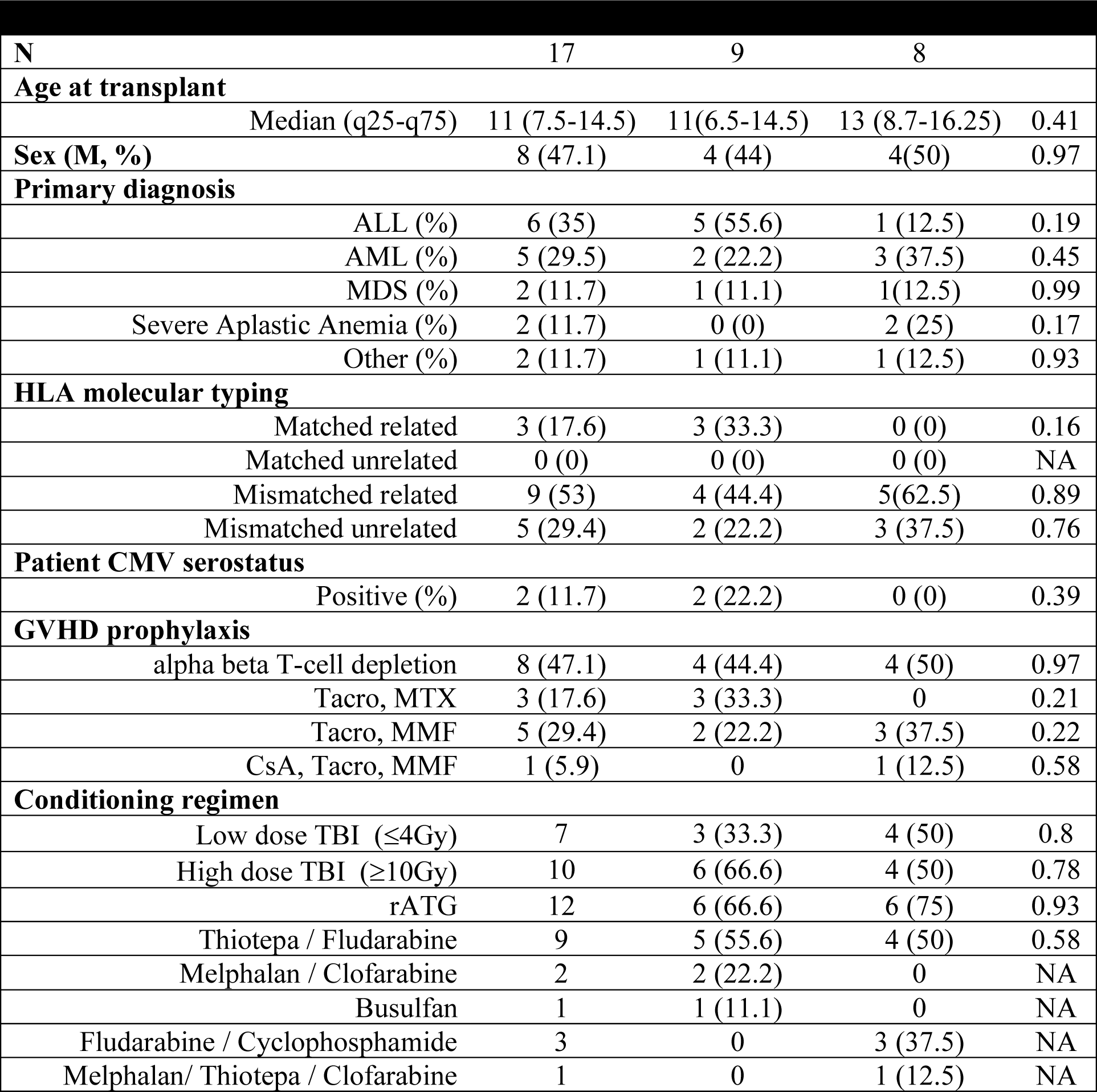
Demographics for the cohort of patients who received HSCT; including primary diagnosis, HLA molecular typing, transplant conditioning regimen and GVHD prophylaxis.

| N | 17 | 9 | 8 |  |
| --- | --- | --- | --- | --- |
| **Age at transplant** |  |  |  |  |
| Median (q25-q75) | 11 (7.5-14.5) | 11(6.5-14.5) | 13 (8.7-16.25) | 0.41 |
| **Sex (M, %)** | 8 (47.1) | 4 (44) | 4(50) | 0.97 |
| **Primary diagnosis** |  |  |  |  |
| ALL (%) | 6 (35) | 5 (55.6) | 1 (12.5) | 0.19 |
| AML (%) | 5 (29.5) | 2 (22.2) | 3 (37.5) | 0.45 |
| MDS (%) | 2 (11.7) | 1 (11.1) | 1(12.5) | 0.99 |
| Severe Aplastic Anemia (%) | 2 (11.7) | 0 (0) | 2 (25) | 0.17 |
| Other (%) | 2 (11.7) | 1 (11.1) | 1 (12.5) | 0.93 |
| **HLA molecular typing** |  |  |  |  |
| Matched related | 3 (17.6) | 3 (33.3) | 0 (0) | 0.16 |
| Matched unrelated | 0 (0) | 0 (0) | 0 (0) | NA |
| Mismatched related | 9 (53) | 4 (44.4) | 5(62.5) | 0.89 |
| Mismatched unrelated | 5 (29.4) | 2 (22.2) | 3 (37.5) | 0.76 |
| **Patient CMV serostatus** |  |  |  |  |
| Positive (%) | 2 (11.7) | 2 (22.2) | 0 (0) | 0.39 |
| **GVHD prophylaxis** |  |  |  |  |
| alpha beta T-cell depletion | 8 (47.1) | 4 (44.4) | 4 (50) | 0.97 |
| Tacro, MTX | 3 (17.6) | 3 (33.3) | 0 | 0.21 |
| Tacro, MMF | 5 (29.4) | 2 (22.2) | 3 (37.5) | 0.22 |
| CsA, Tacro, MMF | 1 (5.9) | 0 | 1 (12.5) | 0.58 |
| **Conditioning regimen** |  |  |  |  |
| Low dose TBI ($\leq 4$Gy) | 7 | 3 (33.3) | 4 (50) | 0.8 |
| High dose TBI ($\geq 10$Gy) | 10 | 6 (66.6) | 4 (50) | 0.78 |
| rATG | 12 | 6 (66.6) | 6 (75) | 0.93 |
| Thiotepa / Fludarabine | 9 | 5 (55.6) | 4 (50) | 0.58 |
| Melphalan / Clofarabine | 2 | 2 (22.2) | 0 | NA |
| Busulfan | 1 | 1 (11.1) | 0 | NA |
| Fludarabine / Cyclophosphamide | 3 | 0 | 3 (37.5) | NA |
| Melphalan/ Thiotepa / Clofarabine | 1 | 0 | 1 (12.5) | NA |

**7).**
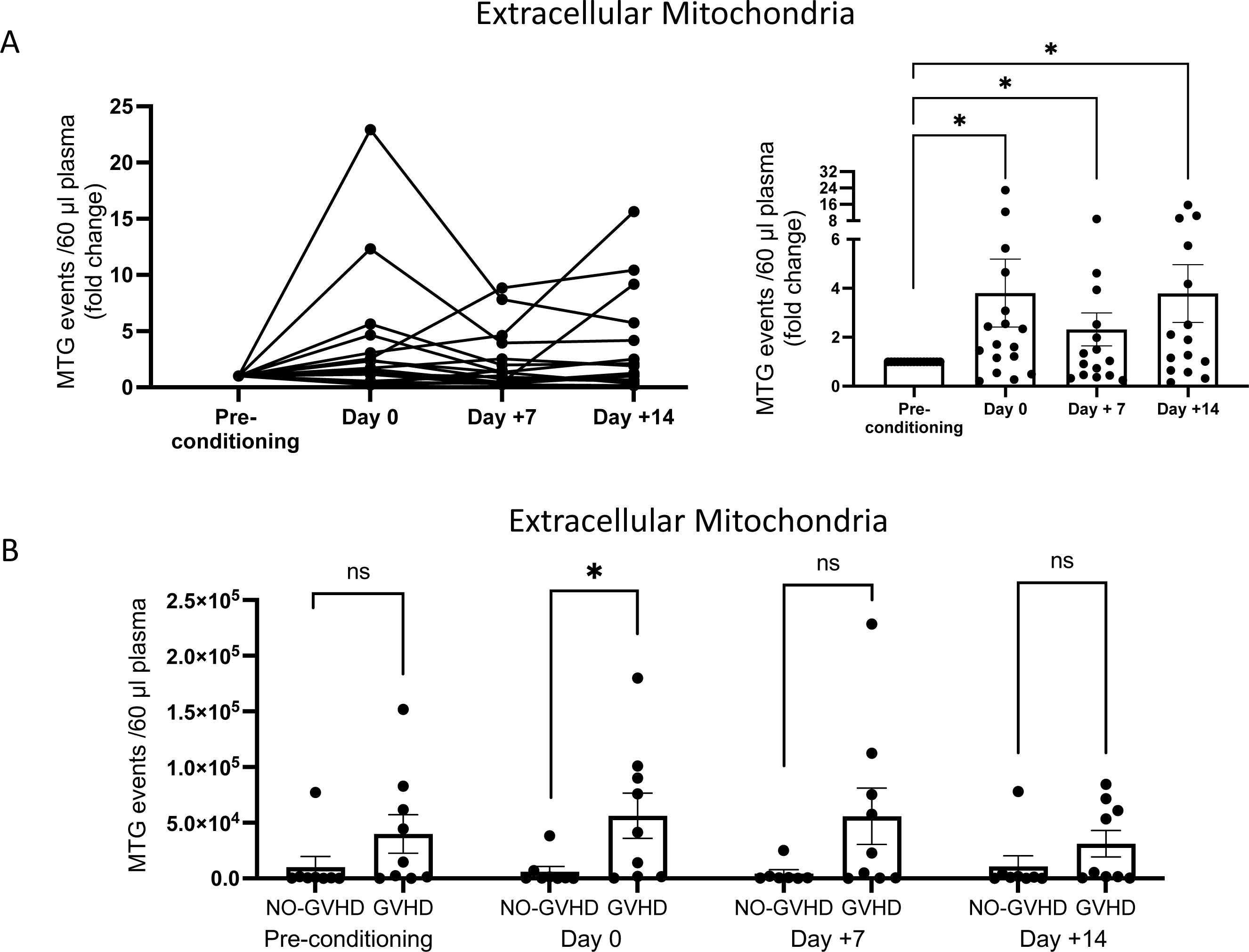
TBI conditioning increases circulating exMito in patients undergoing HSCT. Serial samples collected from HSCT patients prior to initiation of conditioning regimen (pre-conditioning), day of transplant (day 0), days 7 (+7) and 14 (+14) following HSCT was evaluated for abundance of exMito in circulation. **(A)** MTG positive events represented as fold induction relative to the pre-conditioning sample of each patient. **(B)** Comparison of mitochondrial levels in samples collected at the indicated times between patients with and without GVHD. (*p < 0.05)

## 4. Discussion

Although allogeneic T cells have been long understood to be a prerequisite for GVHD, the initiation of GVHD depends on tissue damage, typically resulting from the cytotoxic therapies given as pre-transplant conditioning to ensure HSC engraftment ^(1–4)^. Conditioning-induced damage to mucosa, including the intestine, allows translocation of bacteria which express PAMPs while simultaneously leading to the release of endogenously derived DAMPs ^(6,7)^. Whereas previous studies have linked GVHD severity with the release of these danger signals, the role of exMito in the pathogenesis of GVHD has not been previously characterized. Here we demonstrate that clinically relevant doses of radiation and chemotherapy which are used as pre-HSCT conditioning cause mitochondrial dysfunction and alteration in mitochondrial dynamics, resulting in exMito release both *in vitro* (Figs. 1, 3A) and *in vivo* (Figs. 2, 3D). The presence of circulating exMito exacerbates GVHD-associated mortality (Fig. 4-6). The clinical relevance of these observations is highlighted by the demonstration that TBI-based conditioning increased the level of circulating cell-free mitochondria in patients undergoing HSCT and correlated with the incidence of acute GVHD post-transplant (Fig. 7).

### 4.1 Immunomodulatory role of extracellular released mitochondria

Mitochondria contain byproducts that can induce immune cell activation via multiple mechanisms, including but not limited to, mtDNA-mediated TLR9 signaling, N-formyl peptide-mediated receptor activation, cardiolipin-mediated pro-inflammatory activation and ATP-mediated signaling through purinergic receptors ^(11,29,30)^. Specific to GVHD, Koehn et.al. demonstrated that extracellular ATP exacerbates GVHD-associated mortality in pre-clinical models by impairing the anti-inflammatory effect of donor myeloid-derived suppressor cells (MDSC) ^(31)^. Our results demonstrate that release of mitochondria to the extracellular space can also stimulate host APCs (Fig. 4), resulting in activation and proliferation of donor T cells (Fig. 5).

A wide array of cytokines which are known contributors and/or markers of GVHD ^(21–28)^ were induced upon exMito stimulation (Fig. 4C, Table S1). There was also upregulation of cytokines or chemokines, such as IL-13, CSF-1, and CXCL5, which may have mixed roles in GVHD pathogenesis ^(32–36)^. Some molecules like IL-19 have not been specifically studied in GVHD but are associated with autoimmune diseases like psoriasis ^(37)^ (Fig. 4C, Table S1). Much of our current knowledge on the immune stimulation by mitochondria comes from studying specific mitochondrial byproducts (mtDAMPs) in isolation. With the recent discoveries that intact cell-free mitochondria can be found in circulation ^(38,39)^, future multi-omic and single cell approaches are needed to comprehensively characterize the many potential immunostimulatory mitochondrial molecules which are released in the circulation and their impact on the immune phenotypes under physiologic and pathologic conditions.

### 4.2 Mitochondrial dynamics and extracellular release

Mitochondrial dynamics, the balance between mitochondrial fusion, fission, biogenesis and mitophagy, plays an essential role in mitochondrial quality control ^(40)^. Once damaged, mitochondria are segregated from the network by Drp1-dependent mitochondrial fission and subsequently, removed by mitochondrial autophagy ^(41)^. Similar to our findings, previous studies have demonstrated mitochondrial damage, represented by early impairment in both oxidative phosphorylation and glycolysis shortly after irradiation (Fig. 1 & supplementary Fig. 1)^(42)^, with subsequent increase in glycolysis as a compensatory mechanism. We demonstrate that this impairment in mitochondrial function is coupled with alterations in mitochondrial dynamics, in particular increased mitochondrial fission (Fig. 2). However, we did not observe changes in mitophagy or accumulation of intracellular mitochondrial content following conditioning (Supplemental Fig. S3). Together, this suggests that fragmented mitochondria might be cleared by an alternate mechanism.

Recent publications have suggested that extracellular release of damaged mitochondria is an alternate mechanism by which cells maintain quality control, especially when canonical mechanisms of mitochondrial dynamics are impaired ^(13,43)^. Our data demonstrating an increase in damaged exMito following TBI independent of cell death (Fig. 3), suggest that this mechanism plays an important role in the clearance of damaged mitochondria following conditioning. The underlying mechanism for the extracellular release of mitochondria has been a topic of intense investigation in recent years. Although the consensus is that cell-free mitochondria are a byproduct of intracellular mitochondrial quality control, various mechanisms leading to their generation and release have been proposed ^(43, 44)^. Work from our group has shown that excessive mitochondrial fission is associated with exMito release in pre-clinical models of sepsis and neuroinflammation ^(15,16)^. Independently, other studies have reported that impairments in the classical autophagy pathway contributes to the release of mitochondria ^(13,43)^. Hence, we propose that pharmacologic agents which block excessive mitochondrial fission or induce mitophagy might abrogate the extracellular release of mitochondria and reduce GVHD. Supporting this notion, a recent study reported that baicalin, a flavonoid purified from the plant *Scutellaria baicalensis*, induced autophagy and preserved mitochondrial architecture in the intestinal epithelium and reduced intestinal aGVHD ^(45)^. However, the extracellular release of mitochondria was not explored in this study and requires future investigation.

Although our results are focused on mitochondrial damage and extracellular release in intestinal epithelial cells, it is highly likely that other cell types in various end-organs are sensitive to conditioning mediated mitochondrial damage and therefore contribute to the exMito content in the circulation. This is supported by our observations of impaired bioenergetics and oxidative stress in both HEK cells and HUVECs following irradiation (Supplemental Fig. S11).

In conclusion, the present studies introduce a new pathogenic mechanism for GVHD and suggests potential novel approaches to GVHD prevention, raising several questions that need to be addressed in future investigations. Our results have shown that exMito increase the incidence and severity of acute GVHD. However, the molecular mechanism of exMito-mediated host APC activation require further investigation. Additionally, while our results suggest that both CD4+ and CD8+ T-cells are activated in response to extracellular mitochondria (Fig. 5), further characterization of the T-cell subsets, in both murine and human models would further the understanding of this novel mechanism of GVHD pathogenesis. Finally, our clinical observations need to be validated in a larger cohort of patients to determine the effects of underlying diagnosis and conditioning methods on exMito levels and GVHD.

Current GVHD prophylaxis strategies consist of globally immune suppressive drugs that are prone to complications including opportunistic infections, viral reactivation as well as engraftment failure. Future studies aiming to block the release of mitochondria into the extracellular milieu or scavenge existing exMito from circulation as a prophylaxis before HSCT are warranted to determine whether exMito could serve as a clinically useful target for prevention of GVHD, possibly without risk of immune suppression.

## Author Contribution

V.V., H.Y., K.W. and B.H. contributed to manuscript preparation. V.V., K.W. and B. H. generated the hypothesis. R.S.N., A.B. and D.M-R. aided with experimental design. V.V., H.Y., B.H., J.K.L., K.A.P., A.E., K.P., J.B., conducted the *in vitro* and animal experiments and analysis. V.V., R.V.P., G.B., I.L. collected and conducted the analysis on the clinical samples. N.O. preformed the blinded imaging analysis. All authors edited the manuscript.

---

**S1).**
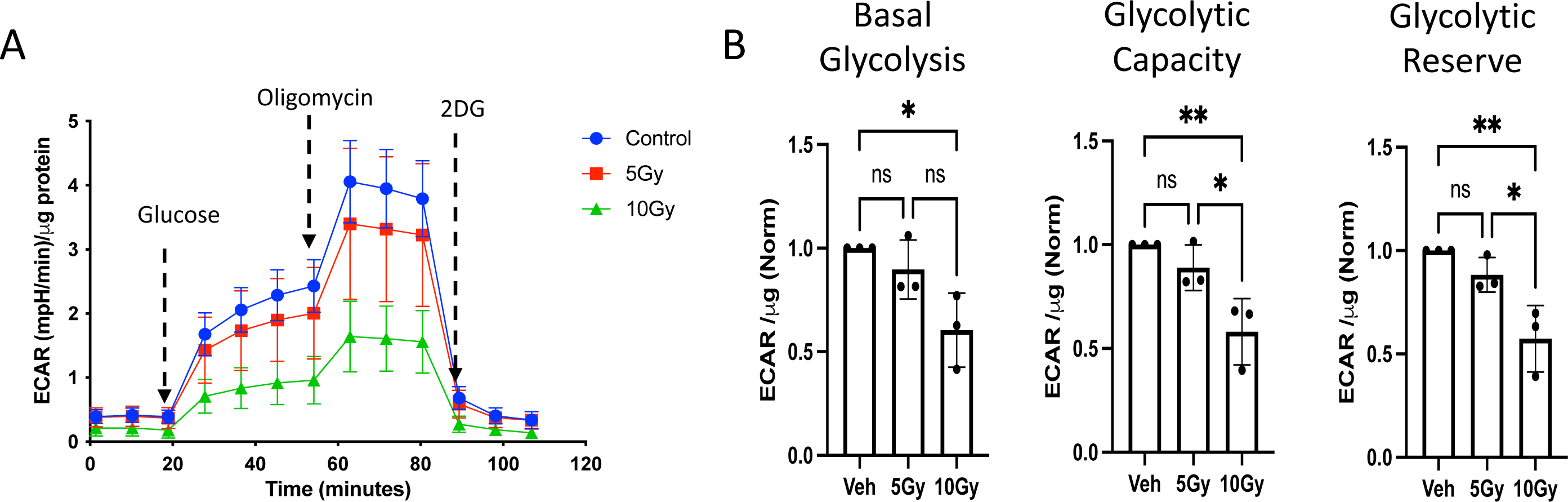
Irradiation mediates impairments in glycolysis in vitro: Caco-2 cells were irradiated at the indicated doses, cultured for an additional 24 hours and subjected to glycolysis stress test using seahorse xfe24 **A)** A representative Seahorse plot of the glycolysis stress test and **(B)** bar graphs of the basal glycolysis, glycolytic capacity and glycolytic reserve normalized to the total protein content (n=3). (*p < 0.05, **p < 0.01; n-number of independent experiments)

**S2).**
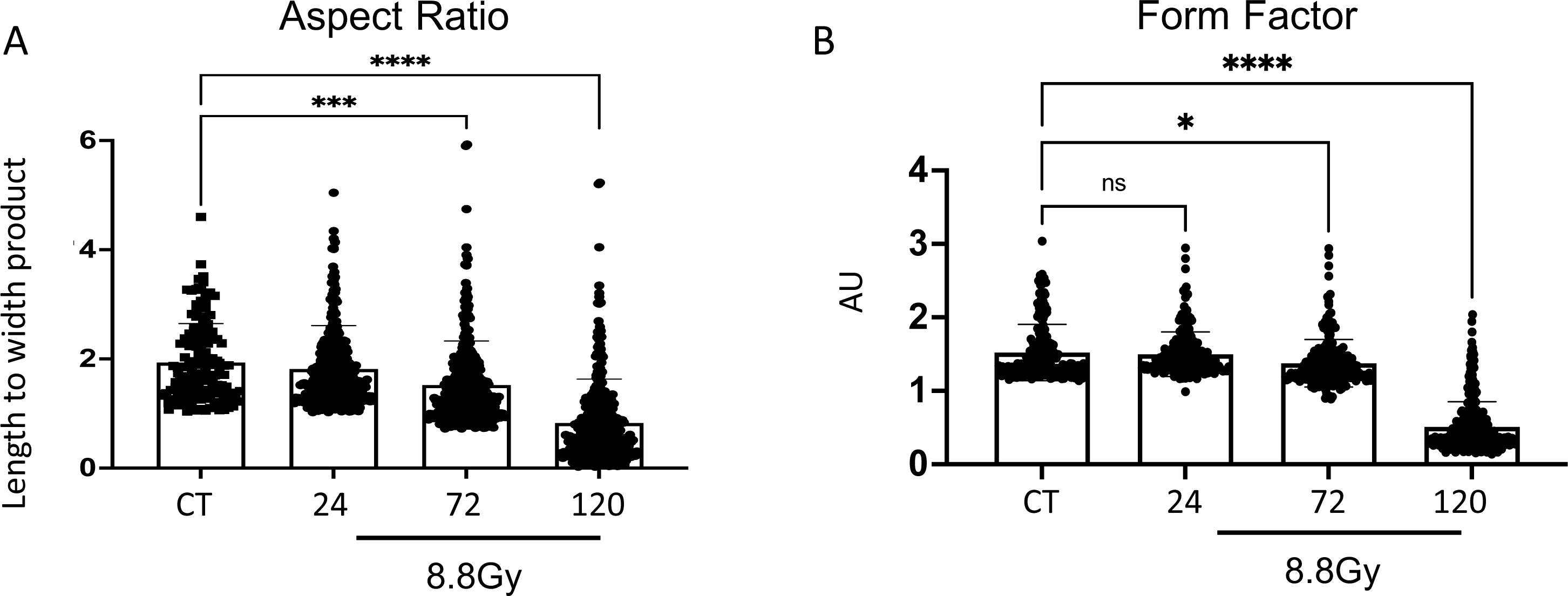
TBI induces mitochondrial damage in the small bowel. Mice were subjected to TBI (8.8 Gy), and the small bowel was harvested at the indicated time points **(A)** Aspect ratio of mitochondria calculated from electron micrographs **(B)** form factor of mitochondria calculated from electron micrographs (*p < 0.05, **p < 0.01, ***p < 0.001, ****p < 0.0001; n-number of independent experiments).

**S3).**
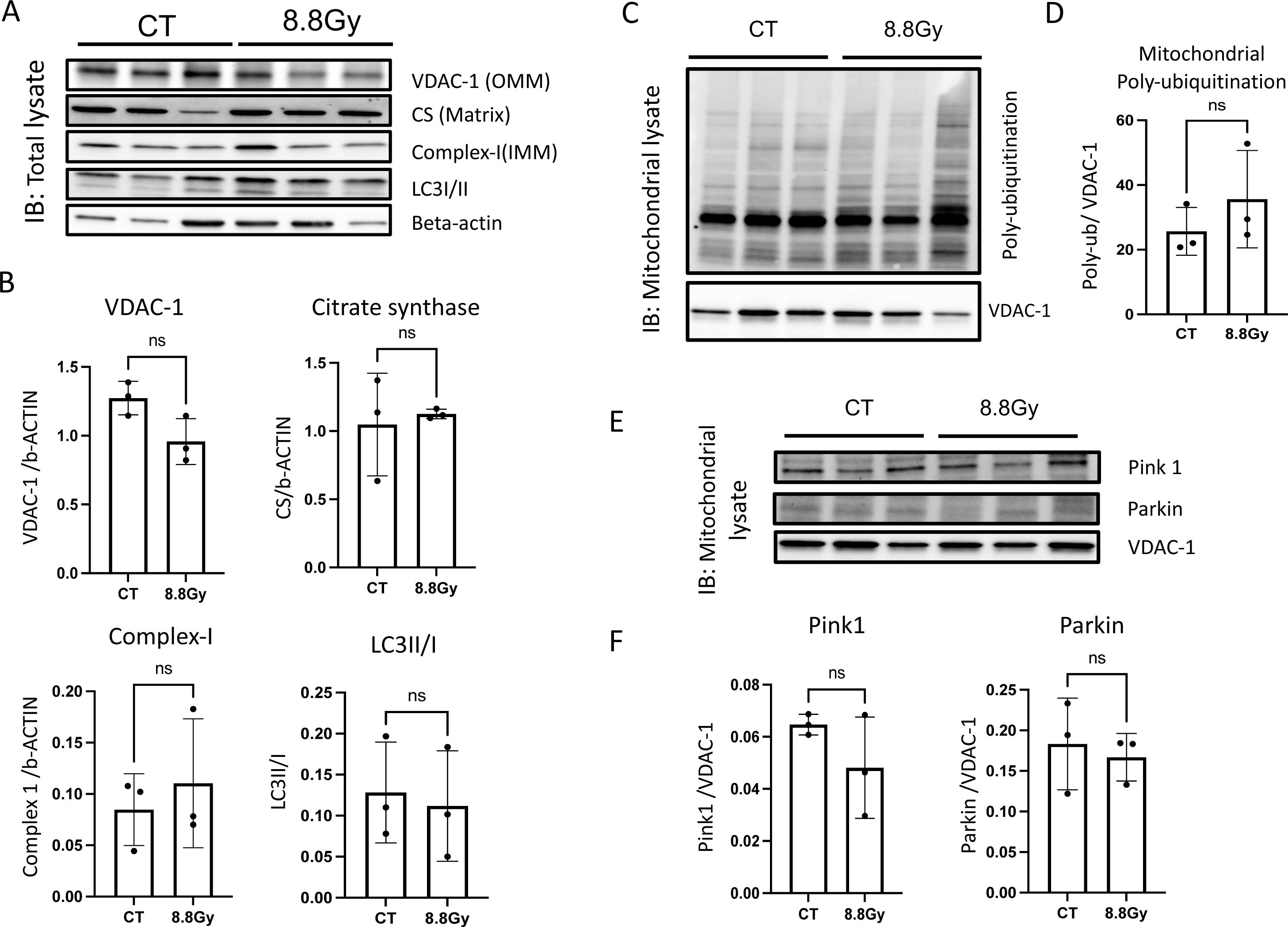
TBI is not associated with intracellular accumulation of mitochondria or induction of mitophagy in the bowel. Representative western blots of total tissue lysates from bowel of mice collected 24 hours following irradiation **(A)** and probed with the indicated antibodies. **(B)** The respective densitometric quantifications are shown. **(C)** Representative Western blot and **(D)** densitometric quantifications of the isolated mitochondrial lysates from bowel of mice 24 hours following irradiation probed with an antibody against ubiquitin and VDAC-1. **(E)** Representative Western blots from isolated mitochondrial lysates collected after 24 hours of irradiation and probed with the indicated antibodies. **(F)** The respective densitometric quantifications are shown. (*p < 0.05, **p < 0.01, ***p < 0.001, ****p < 0.0001; n-number of independent experiments).

**S4).**
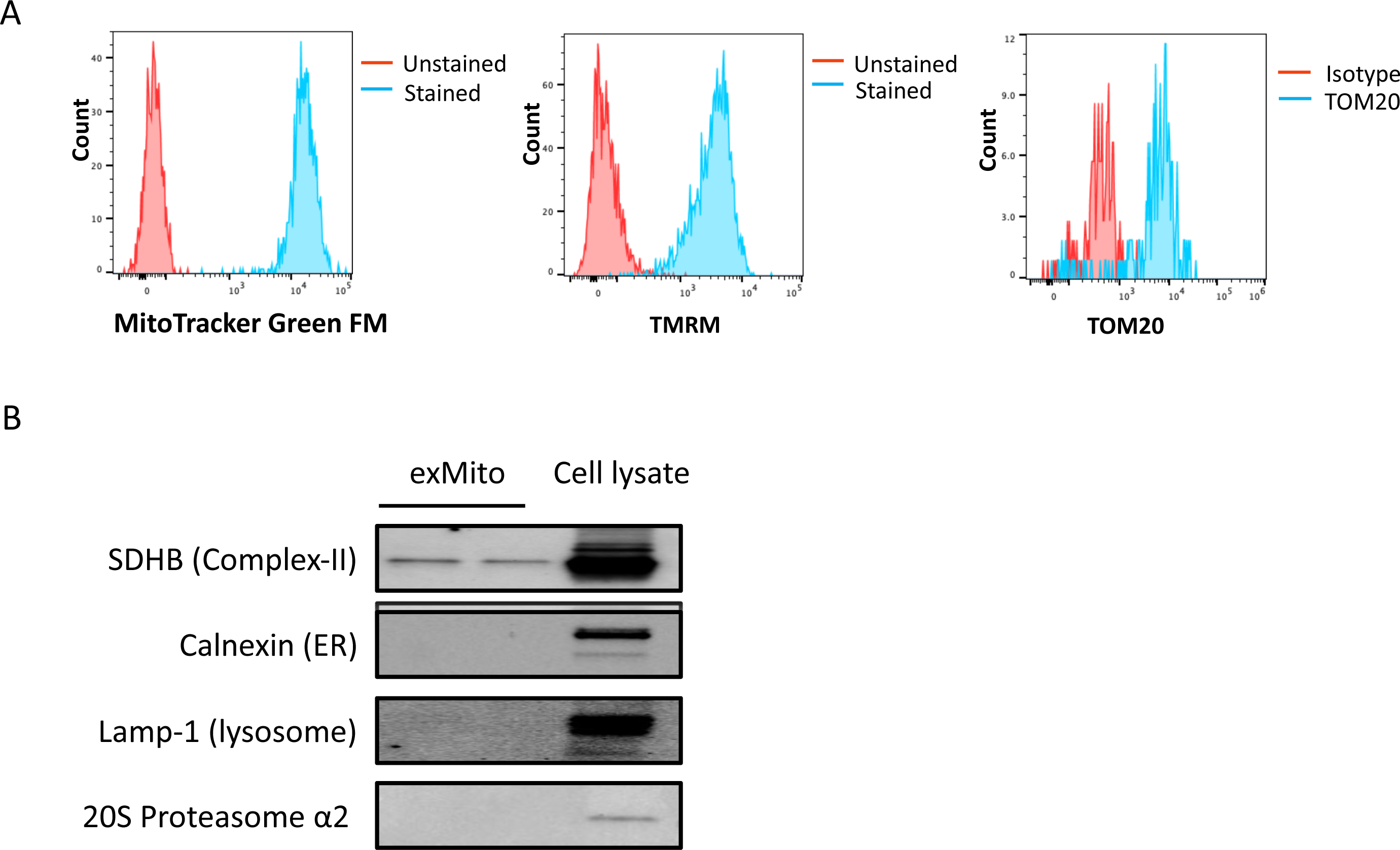
Characterization of Isolated exMito pellet. Isolated pellet from cell culture supernatants of Caco-2 was stained with MTG, TMRM and an antibody for TOM20 and analyzed by flow cytometry. **(A)** A representative histogram for each staining is shown. **(B)** A representative western blot assessing the purity of exMito by evaluating presence of mitochondria associated organelles including Lysosome (Lamp-1) and ER (Calnexin) proteasome (20s proteasome α2).

**S5).**
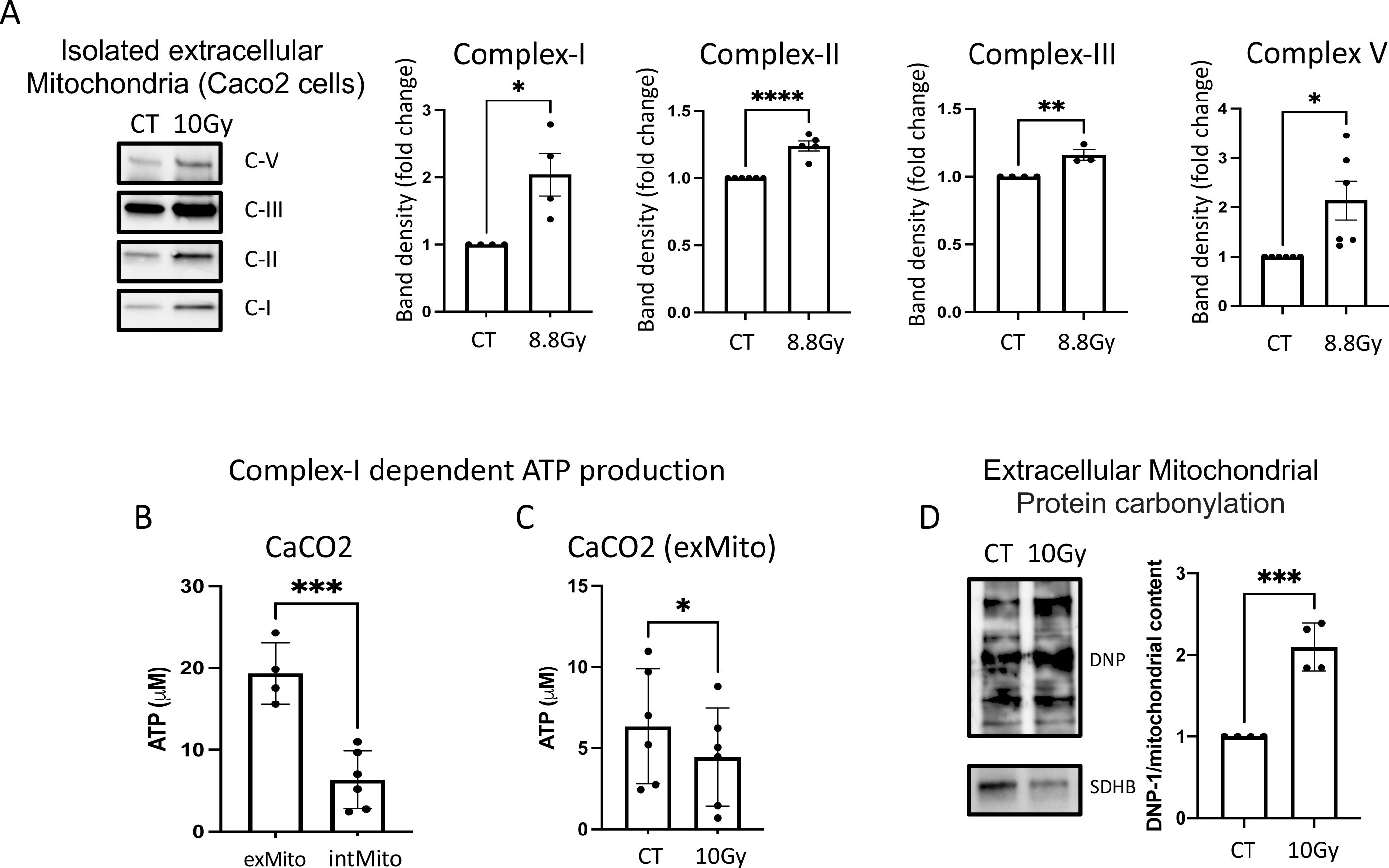
Irradiation mediates release of damaged exMito from Caco-2 cells. **(A)** A representative Western blot and densitometric quantifications of isolated mitochondrial pellet from equal volume of cell culture supernatants of Caco-2 cells 24 hours after treatment with or without irradiation (10Gy) was probed with antibodies against the various electron transport chain complexes. **(B)** The concentration of ATP produced from 1 microgram of isolated intracellular (intMito) and extracellular mitochondria (exMito) when provided with pyruvate and malate is shown (n=4). **(C)** pyruvate and malate-dependent ATP production in isolated exMito from non-irradiated (CT) and irradiated (10Gy) cell culture supernatants (n=6). **(D)** Western blot of isolated exMito from Caco-2 cells probed with antibodies against DNP for protein carbonylation and SDHB for total mitochondrial content (left) and densitometric protein quantification (n=4) (right) is shown. (*p < 0.05, **p < 0.01, ***p < 0.001, ****p < 0.0001; n-number of independent experiments)

**S6).**
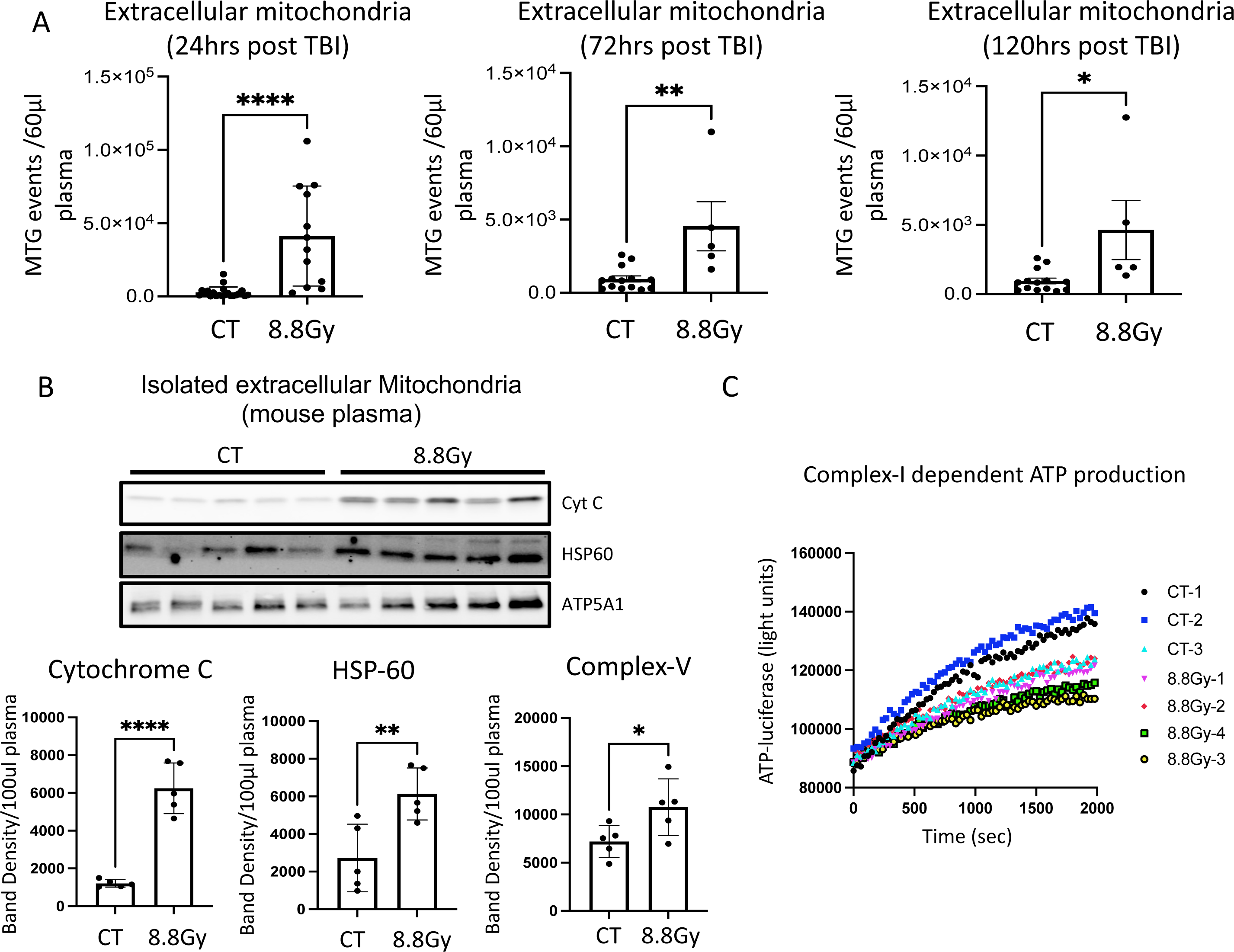
TBI increases exMito in circulation of mice. **(A)** Mouse plasma collected at the indicated times post TBI was stained with MTG was analyzed by flowcytometry for the number of MTG positive events in equal volume of sample. Data is represented as bar graphs (n≥5). **(B)** Representative western blots and their densitometric quantifications of isolated exMito from equal volumes of plasma probed with antibodies against the indicated mitochondrial proteins (n=5). **(C)** A representative plot of luciferase activity recorded over time for ATP production in exMito from plasma is shown. (*p < 0.05, **p < 0.01, ****p < 0.0001; n-number of animals).

**S7).**
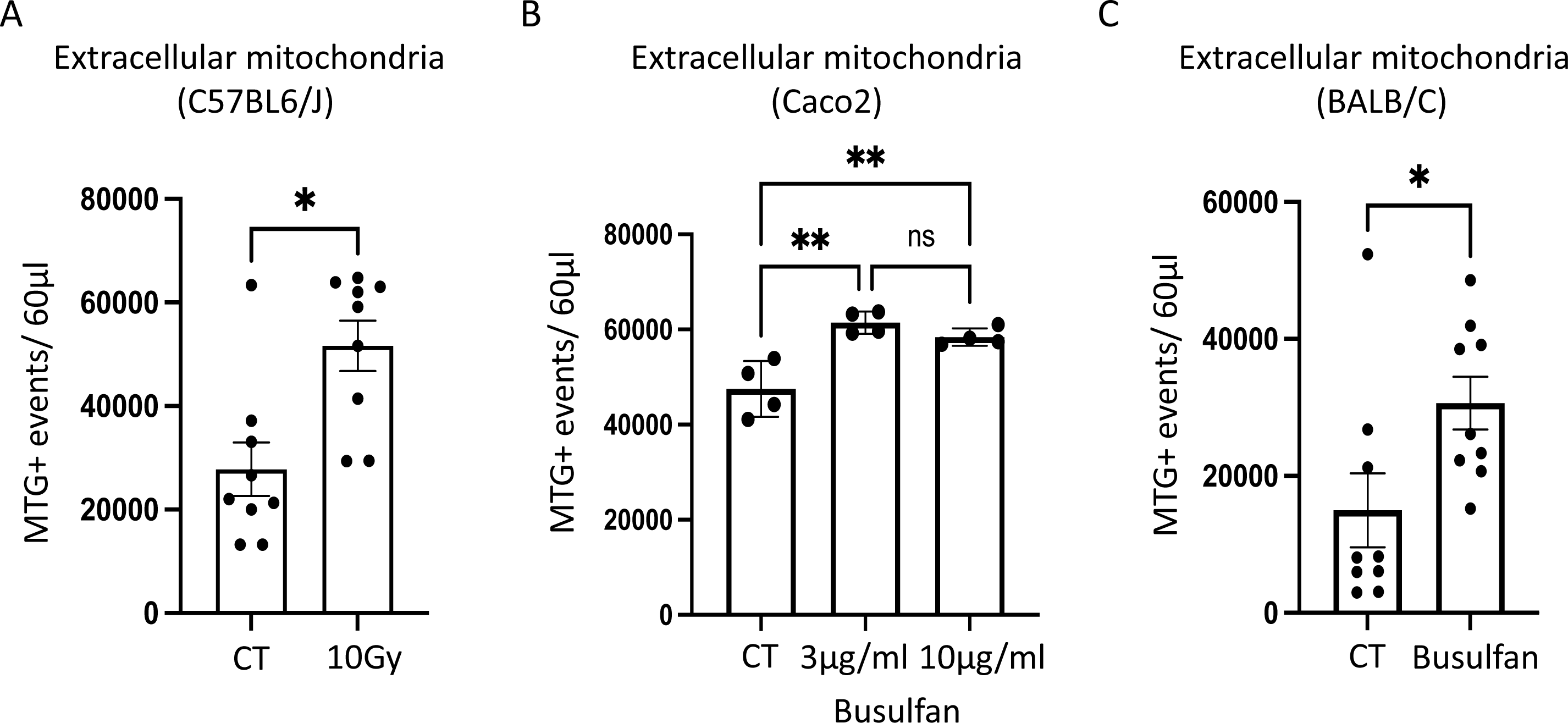
Busulfan conditioning mediates release of exMito similar to TBI: **(A)** exMito content in plasma of C57BL6/J mice 24 hours following TBI (10Gy) was stained with MTG was analyzed by flow cytometry. The number of MTG positive events detected in 60 µL of plasma is shown relative to non-irradiated control mice (n=9). **(B)** exMito content from cell-culture supernatants of control (CT) and Busulfan (3ug/ml and 10ug/ml) treated Caco-2 cells quantified by MTG staining on flow cytometry. **(C)** exMito content in plasma of BALB/c mice 24 hours following myeloablative doses of busulfan (20mg/kg IP x 2 doses) was stained with MTG and analyzed by flow cytometry. The number of MTG positive events detected in 60 µL of plasma is shown relative to non-irradiated control mice (n=10) **(C).** (*p < 0.05, **p < 0.01; n-number of animals).

**S8).**
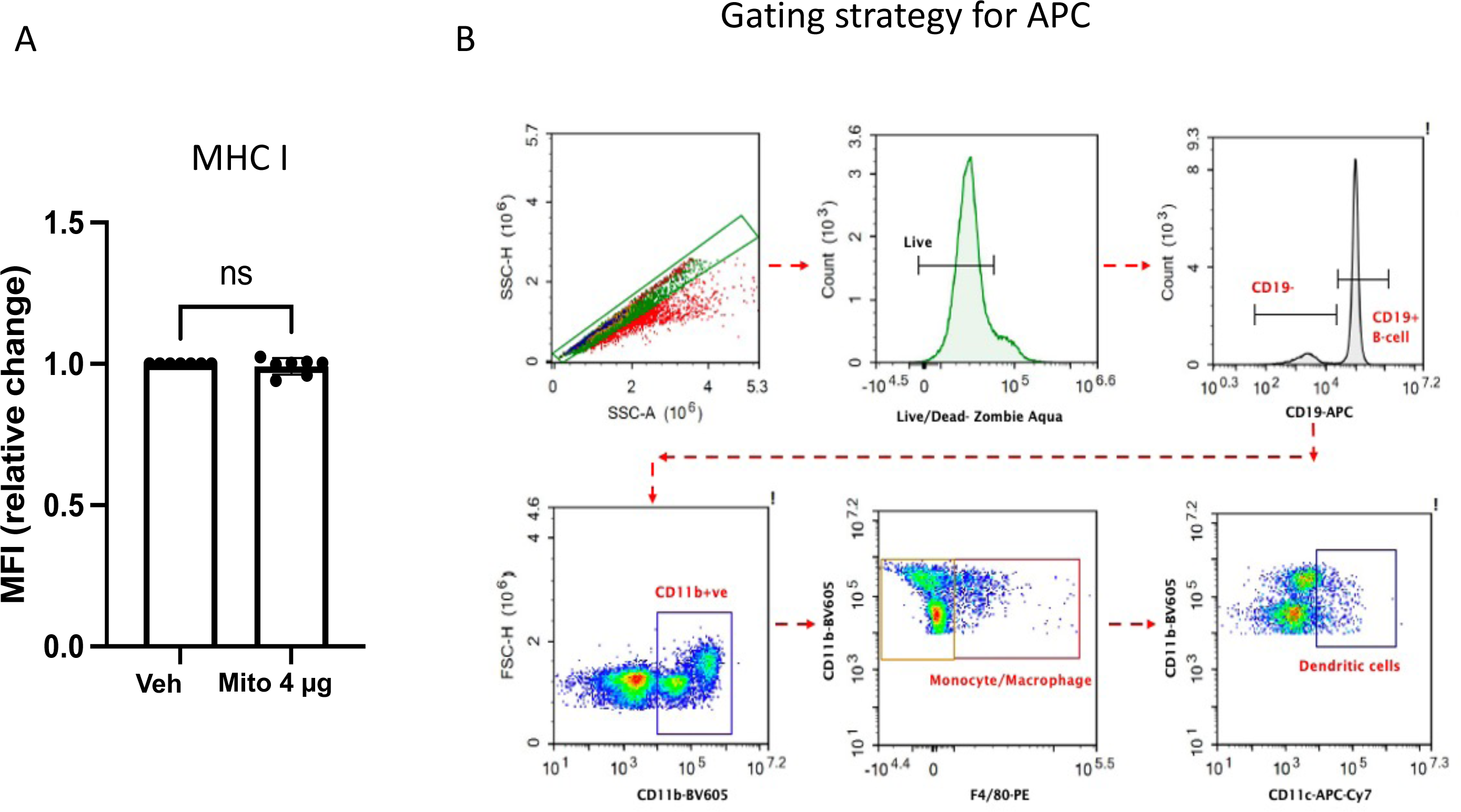
exMito stimulates antigen presenting cells: **A)** T-cell depleted BALB/c splenocytes were treated with 4 µg of mitochondria for 24hours. Cells were stained with an MHC-I antibody and the quantified median fluorescence intensity (MFI) is shown as relative fold change (n=8) **B)** Gating strategy for antigen presenting cell subpopulation including B cell (CD19+), macrophages/monocytes (CD19-, CD11b+, F4/80+) and dendritic cells (CD19-, CD11b+, CD11c+, F4/80-)

**S9).**
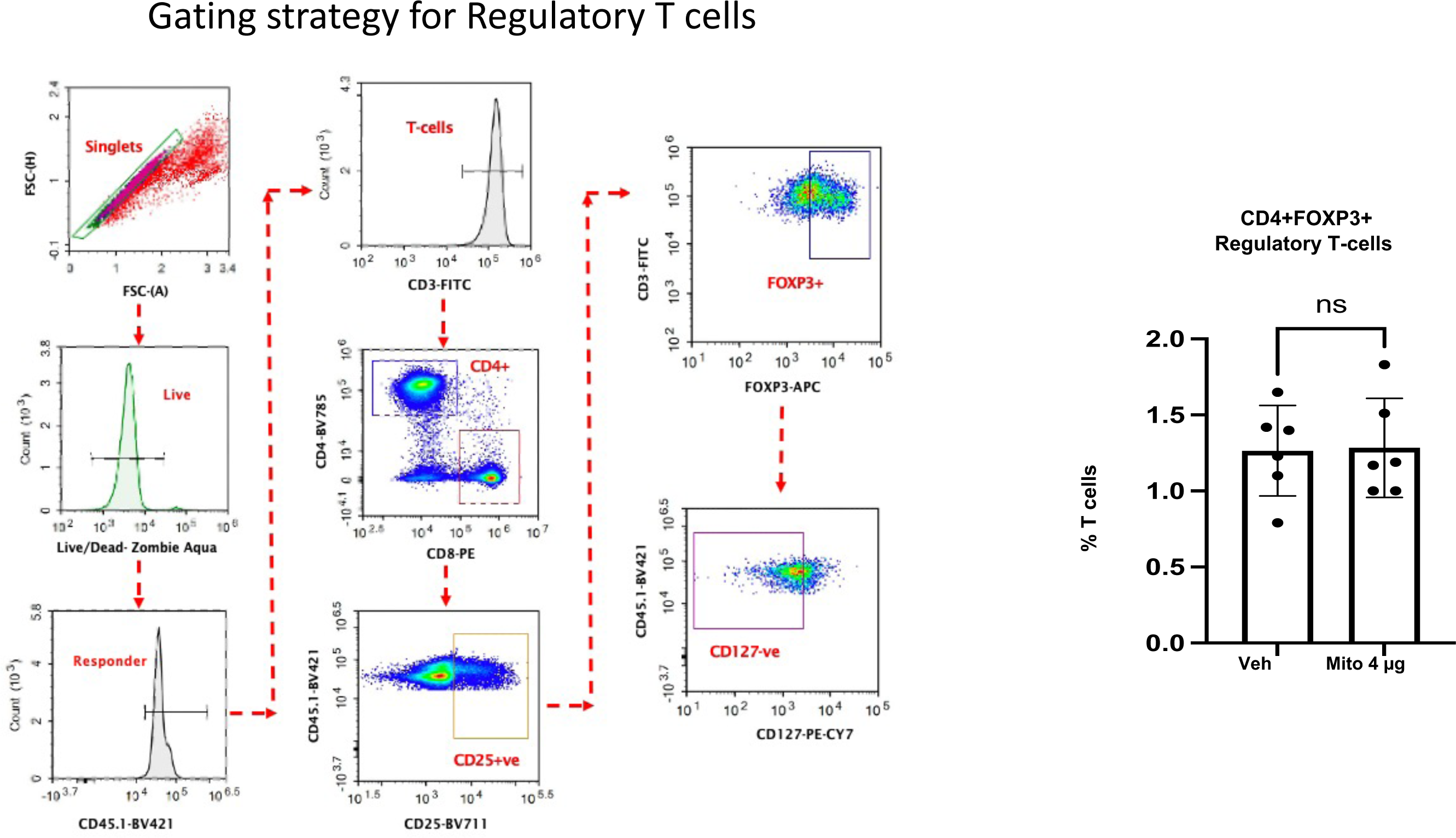
exMito stimulation of antigen presenting cells cause allogenic T-cell activation. *In vitro* MLR between C57BL/6J responder T-cells and BALB/c T-cell depleted splenocytes was set up as described in the Methods section. Gating strategy (left) and percent of regulatory (CD4+, CD25+ FOXP3+ CD127-) T -cells (right) after 48 hours in a MLR (n=6) (n-number of experiments).

**S10).**
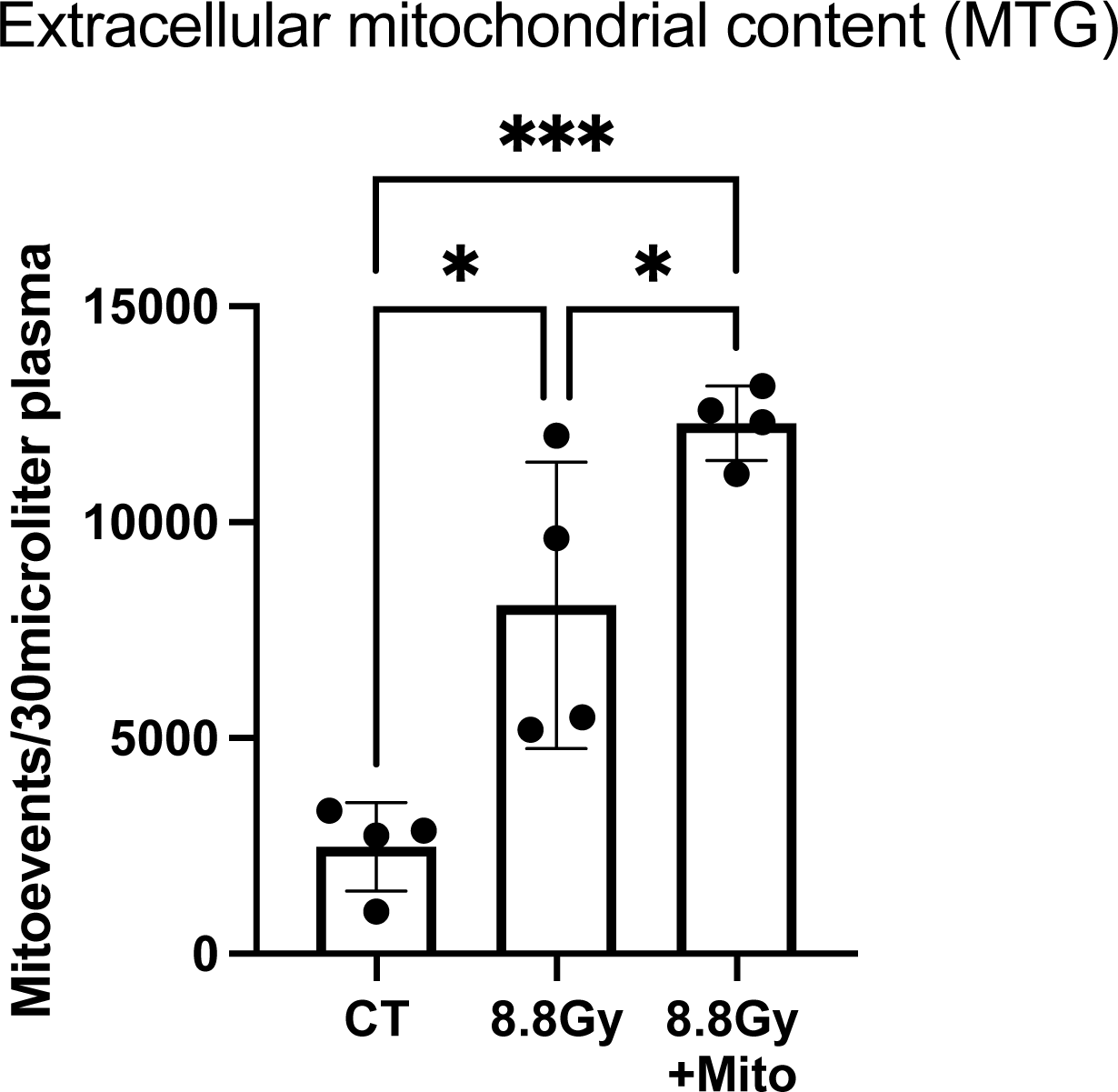
Quantification exMito in circulation following TBI and mito dosing. Quantification of exMito abundance in circulation of mice that were treated with TBI alone or TBI with isolated mitochondria 100ug (n=4). (*p < 0.05, ***p < 0.001, n-number of animals)

**S11).**
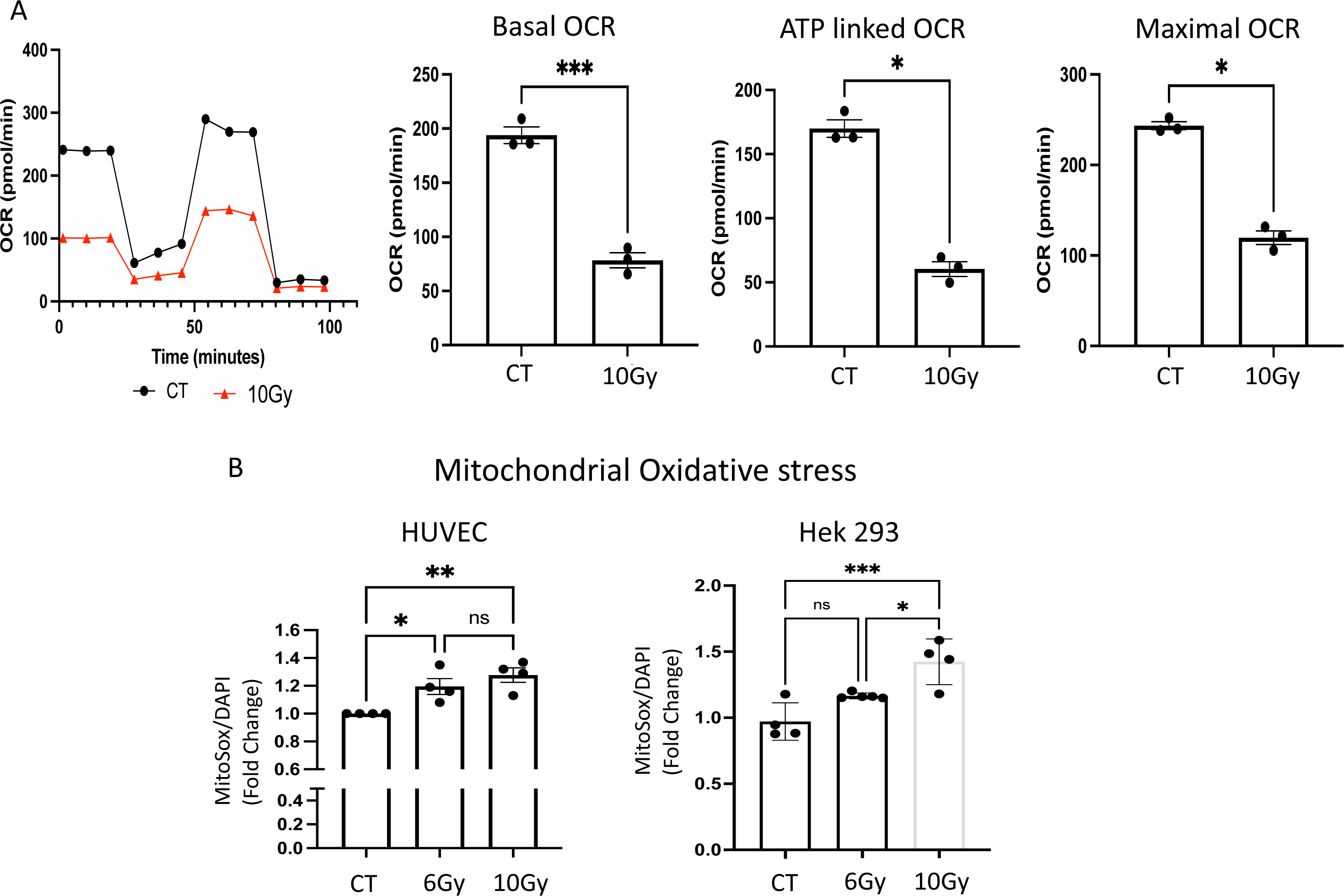
Irradiation impairs mitochondrial function in HEK 293 and HUVEC cell lines and primary endothelial cell cultures. **(A)** HEK293 cells were left untreated (CT) or treated with 10Gy irradiation and after 24 h subjected to Seahorse Mitostress assay. A representative Seahorse plot and bar graphs of the respiratory parameters are shown (n=3). **(B)** HUVECs and HEK293 cells were left untreated (CT) or treated with the indicated doses of irradiation, stained with MitoSOX^TM^ after 24 hours and analyzed by flow cytometry (n≥3). (*p < 0.05, **p < 0.01, ***p < 0.001; n-number of independent experiments).

**Supplemental Table 1:** List of cytokines and chemokines that was differentially regulated between untreated and exMito treated T-cell depleted splenocytes.

| **IL-10** | 128.96 | 3.4782 |
| --- | --- | --- |
| **M-CSF** | 62.22 | 2.9393 |
| **CCL11** | 21.81 | 2.8618 |
| **IL-33** | 81.15 | 2.6098 |
| **G-CSF** | 55.11 | 2.5942 |
| **CXCL5** | 91.57 | 2.5885 |
| **BTC** | 37 | 2.5662 |
| **IFNa** | 57.81 | 2.4852 |
| **IL-13** | 59.17 | 2.4428 |
| **IL-31** | 51.88 | 2.3837 |
| **IL-19** | 86.83 | 2.1973 |
| **IL-17A** | 79.57 | 2.1517 |
| **IL-9** | 30.4 | 2.1019 |
| **IL-7** | 29.13 | 2.0396 |
| **MIP-1a** | 34.11 | 1.9765 |
| **MCP-3** | 33.29 | 1.9493 |
| **sRANKL** | 24.06 | 1.944 |
| **MIP-2** | 63.95 | 1.8978 |
| **IL-25** | 49.41 | 1.8453 |
| **CXCL10** | 25.3 | 1.841 |
| **IFNg** | 115.74 | 1.7441 |
| **IL-2RA** | 38.92 | 1.7183 |
| **IL-5** | 41.59 | 1.6228 |
| **VEGF** | 35.97 | 1.5854 |
| **LEPTIN** | 32.83 | 1.5493 |
| **IL-6** | 90.85 | 1.4854 |
| **CXCL1** | 34.89 | 1.4394 |
| **ST2/IL-33R** | 14.43 | 1.4307 |
| **CCL5** | 37.98 | 1.3646 |
| **IL-27** | 29.64 | 1.32 |
| **GM-CSF** | 17.2 | 1.1939 |
| **TNF-a** | 15.15 | 1.1485 |
| **IL-15/IL-15R** | 47.46 | 1.1206 |
| **IL-28** | 17.62 | 0.924 |
| **MCP-1** | 16.48 | 0.8975 |
| **IL-1a** | 22.03 | 0.8675 |
| **IL-23** | 13.36 | 0.8215 |
| **CHEX-3** | -4 | 0.8 |
| **IL-3** | 50.72 | 0.6734 |
| **IL-4** | 26.76 | 0.5897 |
| **CHEX-4** | -9 | 0.5 |
| **IL-7RA** | 7.19 | 0.4551 |
| **TNFSF13B** | 9.49 | 0.4417 |
| **IL-2** | 5.75 | 0.4068 |
| **IL-1b** | 9.09 | 0.298 |
| **MIP-1b** | 10.41 | 0.2322 |
| **LIF** | -2.33 | 0.1388 |
| **IL-22** | 6.08 | 0.1288 |
| **IL-12P70** | -0.89 | 0.0668 |
| **IL-18** | -0.34 | 0.0143 |
| **CHEX-1** | -7 | 0 |

**Supplemental Table 2:** List of antibodies used for western blot and FACS; including catalogue number supplier and dilution.

| **Western Blot** |  |  |  |
| --- | --- | --- | --- |
| HSP60 | 15282-1-AP | Proteintech | 1:5000 |
| VDAC-1 | Ab14734 | Abcam | 1:1000 |
| SDHB | 10621-1-AP | Proteintech | 1:5000 |
| ATP5A1 | 14676-1-AP | Proteintech | 1:6000 |
| UQCRC2 | 14742-1-AP | Proteintech | 1:4000 |
| PINK-1 | 23274-1-AP | Proteintech | 1:500 |
| Parkin | Sc-32282 | Santacruz | 1:500 |
| Beta-actin | 3700s | Cell Signaling Technology | 1:2000 |
| Ubiquitin | Sc-8017 | Santacruz Biotechnology | 1:500 |
| LC3 A/B | 12741s | Cell Signaling Technology | 1:1000 |
| PGC1-alpha | Sc-517380 | Santacruz Biotechnology | 1:500 |
| DNP | A150-117A | Bethyl Laboratories | 1:1000 |
| cytochrome-c | Ab110325 | Abcam | 1:1000 |
| **Flow cytometry** |  |  |  |
| TOM20 | CL-488-11802 | Proteintech | 1:400 |
| Isotype IgG control | CL488-30000 | Proteintech | 1:400 |
| CD3 | 100204 | Biolegend | 1:200 |
| CD69 | 104512 | Biolegend | 1:100 |
| CD4 | 100552 | Biolegend | 1:100 |
| CD4 | 100447 | Biolegend | 1:100 |
| CD8 | 100708 | Biolegend | 1:100 |
| CD19 | 115512 | Biolegend | 1:100 |
| CD127 | 25-1271-82 | eBioscience | 1:20 |
| CD11c | 117324 | Biolegend | 1:20 |
| CD44 | 103059 | Biolegend | 1:100 |
| CD62L | 104418 | Biolegend | 1:100 |
| CD11b | 101257 | Biolegend | 1:100 |
| CD25 | 102049 | Biolegend | 1:160 |
| CD45.1 | 110732 | Biolegend | 1:100 |
| CD86 | 105016 | Biolegend | 1:100 |
| F4/80 | 12-4801-82 | eBioscience | 1:100 |
| FOXP3 | 17-5773-82 | eBioscience | 1:20 |
| MHC Class-I | 12-5998-81 | eBioscience | 1:100 |
| MHC Class-II | 56-5321-82 | eBioscience | 1:400 |

